# Reduced representation characterization of genetic and epigenetic differentiation to oil pollution in the foundation plant *Spartina alterniflora*

**DOI:** 10.1101/426569

**Authors:** Mariano Alvarez, Marta Robertson, Thomas van Gurp, Niels Wagemaker, Delphine Giraud, Malika L. Ainouche, Armel Salmon, Koen J. F. Verhoeven, Christina L. Richards

## Abstract

Theory predicts that environmental challenges can shape the composition of populations, which is manifest at the molecular level. Previously, we demonstrated that oil pollution affected gene expression patterns and altered genetic variation in natural populations of the foundation salt marsh grass, *Spartina alterniflora.* Here, we used a reduced representation bisulfite sequencing approach, epigenotyping by sequencing (epiGBS), to examine relationships among DNA sequence, DNA methylation, gene expression, and exposure to oil pollution. We documented genetic and methylation differentiation between oil-exposed and unexposed populations, suggesting that the *Deepwater Horizon* oil spill may have selected on genetic variation, and either selected on epigenetic variation or induced particular epigenotypes and expression patterns in exposed compared to unexposed populations. In support of the potential for differential response to the *Deepwater Horizon* oil spill, we demonstrate genotypic differences in response to oil under controlled conditions. Overall, these findings demonstrate genetic variation, epigenetic variation and gene expression are correlated to exposure to oil pollution, which may all contribute to the response to environmental stress.

## Introduction

Organismal interactions and response to environment are governed by molecular mechanisms, which are among the most basic levels of biological organization. Studies across a diversity of organisms have described the association of genetic variation with environmental factors (Andrew *et al.*, 2013; Feder & Mitchell-Olds, 2003). More recently, transcriptomics studies in natural populations have identified gene expression differences that are associated with phenotypic plasticity, genotype-by-environment interactions, and local adaptation, and that some of these differences are only elicited in natural environments (Alvarez, Schrey, & Richards, 2015; Nagano *et al.* 2012, 2019; Nicotra *et al.*, 2010). Hence, gene expression variation, like genetic variation, can translate into trait variation that contributes to organismal performance with important population- and community-level ecological effects (Alvarez *et al.*, 2015; Hughes, Inouye, Johnson, Underwood, & Vellend, 2008; Schoener, 2011; Whitham *et al.*, 2006).

Additional layers of variation, including chromatin modifications, small RNAs, and other non-coding variants, can mediate changes in genotypic expression and phenotype. However, this type of variation is infrequently studied in natural settings (Alvarez *et al.* 2015; Kudoh 2016; Nagano *et al.* 2012, 2019; Richards *et al.*, 2017). DNA and chromatin modifications, such as DNA methylation, can also vary among individuals within populations (Banta & Richards, 2018; Becker & Weigel, 2012; Richards *et al.*, 2017), and contribute to phenotypic variation by modulating the expression of genes (Alvarez *et al*., 2015, 2018), the types of transcripts that genes produce (Maor, Yearim, & Ast, 2015), the movement of mobile elements (Matzke & Mosher, 2014), and the production of structural variants (Underwood *et al.*, 2018; Yelina *et al.*, 2015). At the same time, changes in genetic sequence or gene expression may cause variation in patterns of DNA methylation, creating a bidirectional relationship that varies across the genome (Meng *et al.*, 2016; Niederhuth & Schmitz, 2017; Secco *et al.*, 2015). Patterns of DNA methylation have been correlated to habitat types, exposure to stress, and shifts in species range (Foust *et al.*, 2016; Liebl, Schrey, Andrew, Sheldon, & Griffith, 2015; Liebl, Schrey, Richards, & Martin, 2013; Richards, Schrey, & Pigliucci, 2012; Verhoeven, Jansen, van Dijk, & Biere, 2010; Weyrich, *et al.*, 2016; Xie *et al.*, 2015). However, it is often unclear whether changes in DNA methylation, and correlated changes in gene expression, are simply a downstream consequence of changes in allele frequencies or if they may manifest through other mechanisms.

In 2010, the *Deepwater Horizon (DWH)* oil spill developed into the largest marine oil spill in history (National Commission on the BP *Deepwater Horizon* oil spill, 2011), and became an opportunity to test ecological and evolutionary hypotheses in a diversity of organisms exposed to this recurrent anthropogenic stress (e.g. Alvarez *et al.* 2018; DeLeo *et al.* 2018; Hazen *et al.* 2010; Kimes *et al.* 2013; Kimes, Callaghan, Suflita & Morris, 2014; Robertson, Schrey, Shayter, Moss, & Richards, 2017; Rodriguez-R *et al.* 2015; Whitehead *et al.* 2012). A mixture of crude oil and dispersants made landfall along 1,773 kilometers on the shorelines of Louisiana, Mississippi, Alabama and Florida (Mendelssohn *et al.*, 2012; Michel *et al.*, 2013). Nearly half of the affected habitat was salt marsh, which supplies valuable ecosystem functions such as providing nurseries for birds and fish, and buffering storm and wave action (Day *et al.*, 2007; Mendelssohn *et al.*, 2012; Michel *et al.*, 2013). Gulf of Mexico salt marshes are dominated by the hexaploid foundation plant species *Spartina alterniflora* (2n=6x=62; Marchant, 1968), which is remarkably resilient to a variety of environmental stressors (Bedre, Mangu, Srivastava, Sanchez, & Baisakh, 2016; Cavé-Radet, Salmon, Lima, Ainouche, & El Amrani, 2018; Pennings & Bertness, 2001; Silliman *et al.*, 2012). Crude oil exposure from the *DWH* oil spill resulted in reduced carbon fixation, reduced transpiration, and extensive above-ground dieback in *S. alterniflora* populations (Lin & Mendelssohn, 2012; Silliman *et al.*, 2012), but oil-affected populations showed partial to complete recovery within seven months of the spill (Lin *et al.*, 2016). However, the genomic and population level mechanisms that underlie this remarkable recovery have been poorly characterized.

In previous studies, we found that in *S. alterniflora* exposed to the *DWH* oil spill, pollution tolerance was correlated to changes in expression of a diverse set of genes, including epigenetic regulators and chromatin modification genes, such as a homolog of SUVH5 (Alvarez *et al.*, 2018). Although *S. alterniflora* populations were partially resilient to the *DWH* spill (Lin & Mendelssohn, 2012), we found evidence of genetic differentiation between individuals from oil-exposed areas and nearby uncontaminated areas (Robertson *et al.*, 2017). We expected that DNA methylation patterns would be divergent between oil exposed and unexposed populations, which might be induced by the environment or result from the genetic differences between exposed and unexposed populations. However, while a few DNA methylation loci (measured via methylation sensitive amplified fragment length polymorphism; MS-AFLP) were correlated with oil exposure, we did not find genome-wide patterns in DNA methylation correlated with oil exposure in *S. alterniflora* (Robertson *et al.*, 2017).

In this study, we used a recently developed reduced representation bisulfite sequencing (RRBS) technique, epigenotyping by sequencing (epiGBS), to generate a more robust DNA sequence and DNA methylation data set (van Gurp *et al.*, 2016). We expected that the increased resolution, both in number and in detail of the markers, provided by this sequencing approach would confirm our previously observed patterns of genetic differentiation, and allow us to identify fine scale DNA methylation structure that was not apparent in our previous study. By aligning our fragments to the *S. alterniflora* transcriptomes (Boutte *et al.*, 2016; Ferreira de Carvalho *et al.*, 2013, 2017) and *Oryza sativa* genome (Kawahara *et al.*, 2013), we expected to assess the relationship between DNA methylation and previously reported gene expression. We predicted that we would find evidence that DNA methylation was correlated with changes in gene expression since some fragments might overlap with the coding regions of genes (Niederhuth & Bewick *et al.*, 2016). In addition, we examined the potential for response to selection by crude oil exposure among genotypes collected from the field in a common garden greenhouse experiment. We predicted that we would find variation in response to crude oil among genotypes, which would indicate existing standing variation in wild populations of *S. alterniflora* that could be acted upon by selection.

## Materials and Methods

### Sample Collection

We collected individuals from the leading edge of the marsh at three contaminated and three neighboring uncontaminated sites near Grand Isle, Louisiana and Bay St. Louis, Mississippi in August 2010, four months after the *DWH* oil spill as described in previous studies (Table 1; Alvarez *et al.*, 2018; Robertson *et al.*, 2017). These sites were naturally variable in conditions, but all sites supported monocultures of *S. alterniflora.* Contaminated sites were identified by the visual presence of oil on the sediment and substantial above-ground dieback of *S. alterniflora* on the leading edge of the marsh with *S. alterniflora* plants growing through the dead wrack. Nearby uncontaminated sites did not have any visible signs of the presence of oil or noticeable dieback of the above ground portions of *S. alterniflora.* Contamination status was later confirmed via National Resource Damage Assessment databases (Robertson *et al.*, 2017). To standardize age and minimize developmental bias in sampling, we collected the third fully expanded leaf from each of eight individuals, spaced 10 meters apart at each of the six sites (N=48). Leaf samples were immediately frozen in liquid nitrogen to prevent degradation, and kept frozen during transport to the University of South Florida for processing and analysis.

**Table 1.** Sampling information across all sites after filtering. Site information includes location and oil status at each site (exposure).

| Site Location | Site Code | Exposure | No. of individuals |
| --- | --- | --- | --- |
| Grand Isle, LA | GIN1 | None | 8 |
| Grand Isle, LA | GIN2 | None | 6 |
| Barataria Bay, LA | GIO1 | Heavily Oiled | 3 |
| Barataria Bay, LA | GIO2 | Heavily Oiled | 7 |
| Bay St. Louis, MS | MSN | None | 6 |
| Bay St. Louis, MS | MSO | Heavily Oiled | 8 |

### DNA extractions and library prep

We isolated DNA from each field-collected sample (N=48) using the Qiagen DNeasy plant mini kit according to the manufacturer’s protocol. We prepared epiGBS libraries *sensu* van Gurp *et al.* (2016). Briefly, isolated DNA was digested with the enzyme PstI, which is sensitive to CHG methylation and biases resulting libraries toward coding regions (van Gurp *et al.*, 2016). After digestion, adapters containing methylated cytosines and variable barcodes were ligated to either end of the resulting fragments. We used the Zymo EZ Lightning methylation kit to bisulfite treat and clean the DNA. Libraries were then amplified with the KAPA Uracil Hotstart Ready Mix with the following PCR conditions: an initial denaturation step at 98°C for 1 min followed by 16 cycles of 98°C for 15 s, 60°C for 30s, and 72°C for 30s, with a final extension of 72°C for 5 min. We used rapid run–mode paired-end sequencing on an Illumina HiSeq2500 sequencer using the HiSeq v4 reagents and the HiSeq Control software (v2.2.38), which optimizes the sequencing of low-diversity libraries (van Gurp *et al.*, 2016).

### Data pre-processing and mapping to transcriptome

We used the epiGBS pipeline (van Gurp *et al.*, 2016) to demultiplex samples, trim adapter sequences, assemble the *de novo* reference sequence, and call single nucleotide polymorphisms (SNPs) and DNA methylation polymorphisms (DMPs) (https://github.com/thomasvangurp/epiGBS). Sequencing depth varied substantially between samples, which we evaluated with a principal components analysis (PCA) on sampling depths across loci. We assumed that an approximately even spread of the samples across PC1 and PC2 with no association of population or oil exposure, would indicate that sampling depth did not bias our downstream analyses (Figure S1). SNPs (the resulting snps.vcf file) and DMPs (methylation.bed) were filtered separately for each individual to include only loci that were sequenced a minimum of ten times (10x depth of coverage), while loci below this coverage were considered missing data. We first removed 10 individuals with high amounts of missing data (>80%), leaving 38 samples across all 6 populations (Table 1). We then retained only loci that were present in more than 50% of individuals, with no more than 70% missing from any one individual (Figure S2). During the course of this filtering, missing data were imputed via a k-nearest neighbors approach (impute, Hastie, Tibshirani, Narasimhan, & Chu, 2018). We also performed genome-wide analyses (redundancy analyses, explained below) a second time with stricter filtering parameters (no more than 50% missing data in any individual, and no more than 20% missing data at each locus, leaving 34 individuals) and obtained nearly identical P-values and F-statistics, although percent variance explained was reduced (Supplementary File 1).

All fragments were mapped to the published *S. alterniflora* transcriptome (Boutte *et al.*, 2016) and the *O. sativa* genome (Michigan State University version 7, Kawahara *et al.*, 2013) using BLAST (Altschul *et al.*, 1997). We used BLAST (Altschul *et al.*, 1997) and RepeatExplorer (Novak, Neumann, Pech, Steinhaisl, & Macas, 2013) to compare our sequenced fragments to the *S. alterniflora* transcriptome (Boutte *et al.*, 2016; Ferreira de Carvalho *et al.*, 2013, 2017) and known repeat elements, respectively.

### Population genetics

All statistical analyses were performed in R v 3.5.3 (R Core Team, 2017). The epiGBS technique, and the sequencing design that we chose, did not provide sufficient sequencing depth to estimate hexaploid genotype likelihoods with confidence, particularly considering the lack of a high-quality reference genome (Boutte *et al.*, 2016; Dufresne *et al.*, 2014). We therefore used the frequency of the most common allele within an individual at each polymorphic locus as a substitute for genotype at each locus. Although this method ignores the various types of partial heterozygosity that are possible in hexaploid *S. alterniflora*, methods do not currently exist for accurate estimation of heterozygosity in polyploids, and the majority of standard population genetic inference assumes diploidy. We assumed that the frequency of the most common allele was likely to underestimate diversity and therefore underestimate divergence between populations, making our tests of differentiation conservative (Meirmans, Liu, & van Tienderen, 2018).

We obtained pairwise F_ST_ values between populations to test for significant differentiation (StAMPP, Pembleton, Cogan, & Forster, 2013). We also used distance-based redundancy analysis (RDA function in the Vegan package v. 2.5-2; Oksanen *et al.* 2017) to minimize false positives (Meirmans, 2015) in assessing isolation by distance using the formula (genetic distance ~ latitude * longitude). We visualized data using principal components analysis (PCA; Figure 1A).

**Figure 1.**
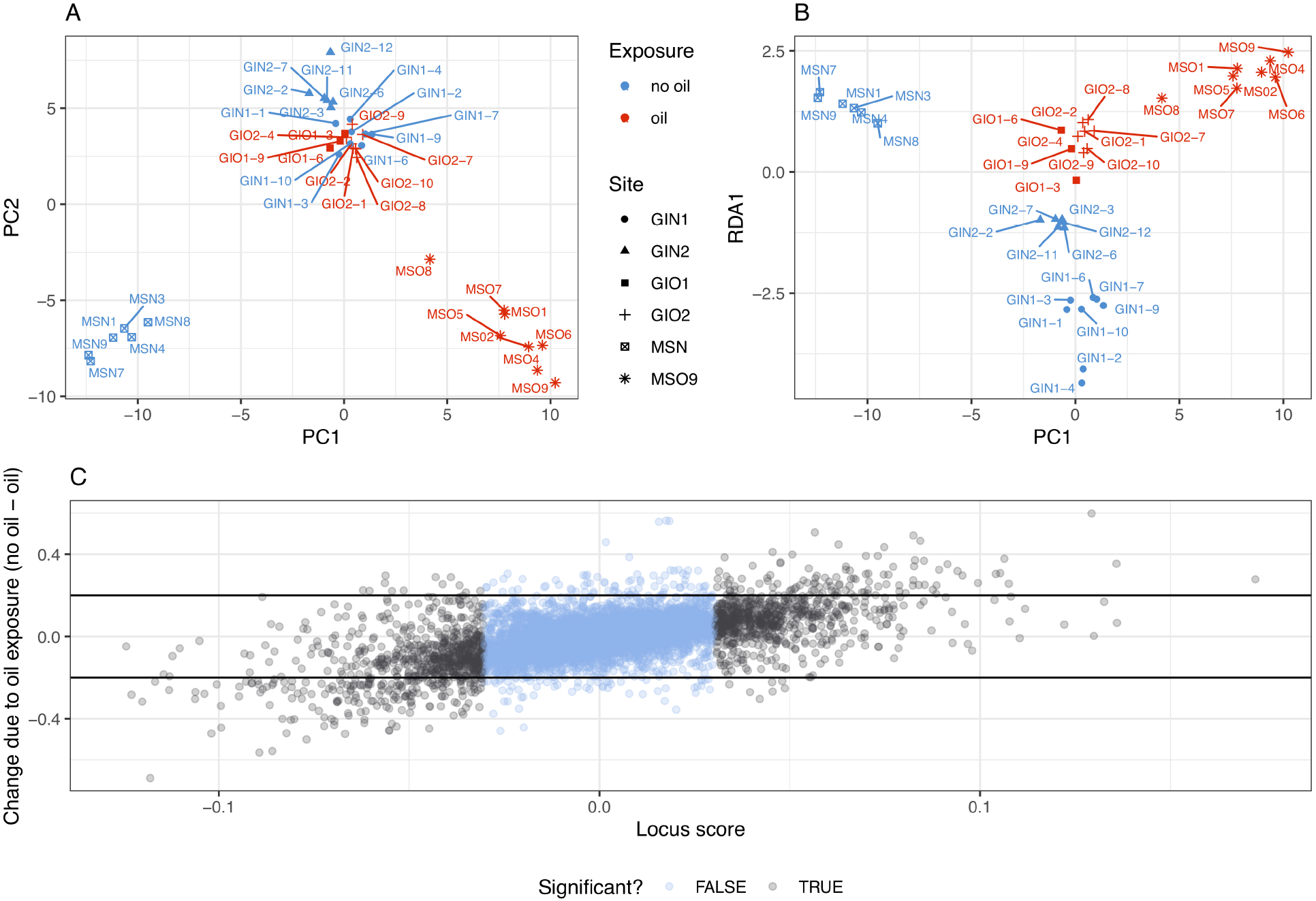
A) Visualization of principal components 1 and 2 of allele frequency data (SNP) data. B) Visualization of principal component 1 and RDA1, which represents the line of maximal separation between samples based on allele frequency data. C) The locus score, representing loadings of each SNP to the constrained axis, plotted against the average change in allele frequency between unexposed and oil exposed populations. Significantly differentiated SNPs are shown in black.

To quantify the relationship between genome-wide variation and environmental conditions, we used partial constrained redundancy analysis (RDA, implemented with the RDA function in the Vegan package v. 2.5-2; Oksanen *et al.* 2017). RDA is a multivariate ordination technique that allowed us to assess the joint influence of all SNPs simultaneously, while effectively controlling for both population structure and false discovery (Forester, Lasky, Wagner, & Urban, 2018). The resulting “locus scores” correspond to the loadings of each SNP on to the constrained axis, which represents the variation that can be explained by the variable of interest (in this case, crude oil exposure). We attempted to control for variation among sites with a replicated sampling strategy, but rather than using a single term for “population”, we conditioned our ordination on variables identified by latent factor mixed models analysis using the LFMM package (Caye, Jumentier, Lepeule, & François, 2019), which provides a method to account for residual variation due to unmeasured differences among populations, including environmental variation, life history variation, and geographical separation (Leek *et al.*, 2017). We used RDA to fit our final model with the formula (SNP matrix ~ oil exposure + Condition(latent factors)). We used a permutational test (999 permutations; Oksanen *et al.* 2017) to assess the likelihood that oil-exposed and unexposed populations differed by chance, and visualized results using principal components analysis. We identified individual SNPs that were significantly correlated with oil contamination using the three standard deviation outlier method described by Forester *et al.* (2018). Finally, we tested for differences in genetic variation using the PERMDISP2 procedure (Vegan; Oksanen *et al.*, 2017), under the assumption that a significant reduction in genetic variation in oiled populations may be evidence of a bottleneck.

### Methylation analysis

During the filtering process, loci were annotated with their methylation context, but all contexts were pooled for distance-based analyses as well as multiple testing correction after locus-by-locus modeling. We tabulated methylation frequency at each locus (methylated cytosines/(methylated+unmethylated cytosines)), and visualized differences between samples with PCA (Figure 2A).

**Figure 2.**
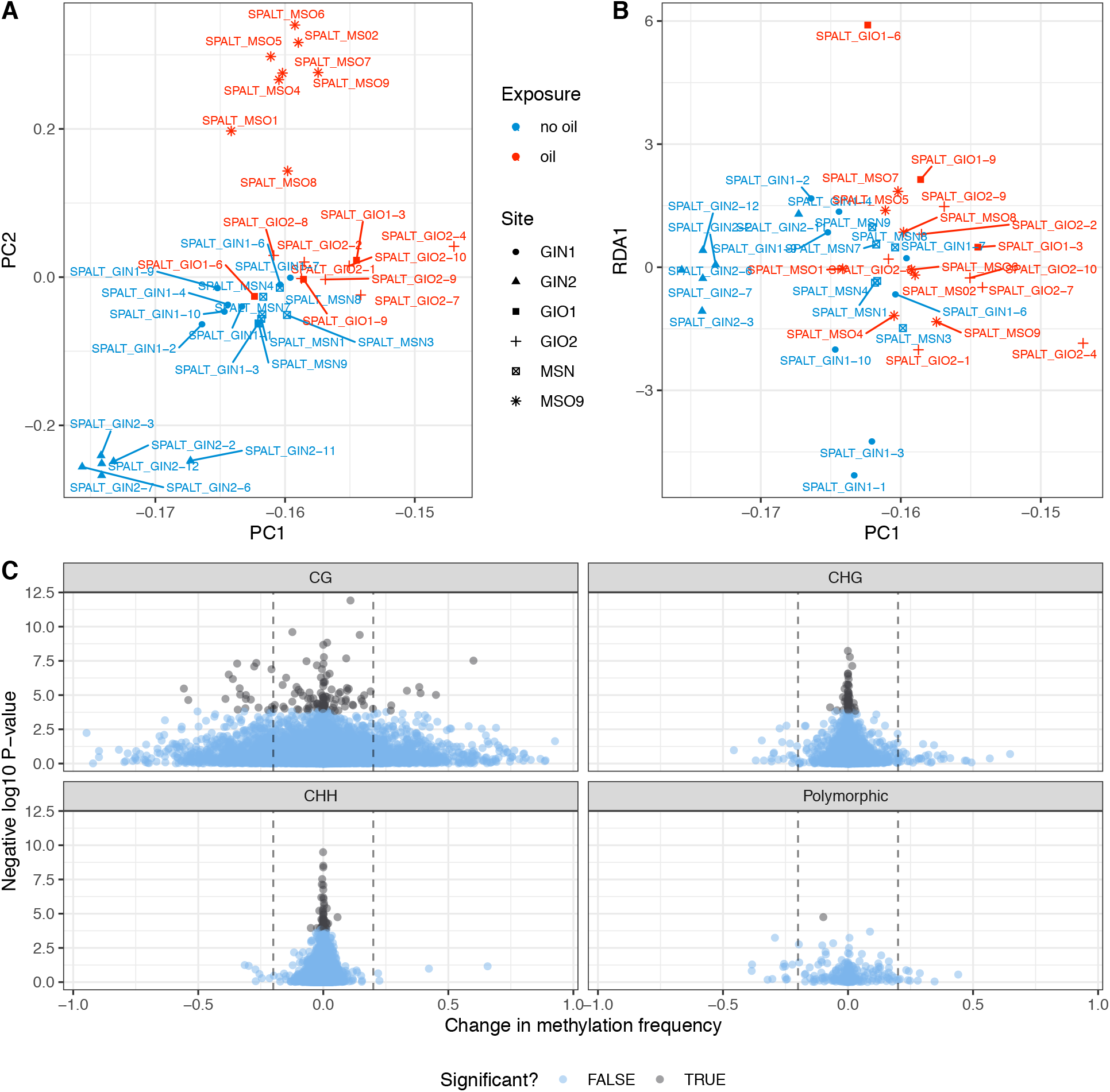
A) Visualization of principal components 1 and 2 of methylation frequency data. B) Visualization of principal component 1 and RDA1, which represents the line of maximal separation between samples based on methylation frequency data, after controlling for genetic variation and latent factors. C) All methylation polymorphisms, with differentially methylated positions (DMPs) shown in black. Negative log10 P-values are plotted on the Y axis while the average change in methylation frequency between unexposed and oil exposed populations is shown on the X axis. Dotted lines represent a change of at least 20%, either increased or decreased, due to oil exposure.

To identify signatures of DNA methylation variation that were correlated with oil exposure while controlling for genetic structure, we estimated latent variables with LFMM (Caye *et al.*, 2019) as above. In addition to the advantages described above, latent factor analysis (or the related surrogate variable analysis) provides a control for cell type heterogeneity in epigenomic studies (Akulenko, Merl, & Helms, 2016; Caye *et al.*, 2019; McGregor *et al.*, 2016). We then modeled the impact of oil exposure to genome-wide patterns of DNA methylation while controlling for latent variation as well as population structure via RDA (Vegan v. 2.5-2; Oksanen *et al.*, 2017) with the formula (methylation distance ~ oil exposure + Condition(latent factors) + the first 5 principal components).

To identify differentially methylated positions (DMPs) between contaminated and uncontaminated samples, we used binomial linear mixed modeling (MACAU; Lea, Tung, & Zhou, 2015), using the genetic relatedness matrix and latent factors as covariates to control for population structure. We corrected locus-specific P-values for multiple testing (qvalue v 2.14.1; Storey, Bass, Dabney, & Robinson, 2015), and tested for overrepresentation of cytosine contexts (CG, CHG, CHH) using binomial tests, implemented in R (R Core Team, 2017). Our epiGBS fragments rarely exceeded 200bp, and we were therefore unable to identify differentially methylated regions.

### Relationships to gene expression variation

In a previous study using pools of individuals on a custom microarray, we found differential expression associated with response to the *DWH* in 3,334 out of 15,950 genes that were assessed in *S. alterniflora* (Alvarez *et al.*, 2018). In order to make the epiGBS data comparable to our pooled microarray design, we concatenated SNPs and methylation polymorphisms from individuals into *in silico* sample pools by averaging values at individual loci across the same three individuals within pools that were used in the gene expression analysis. We then calculated genetic, expression, and methylation distances between sample pools and used Mantel and partial Mantel tests to assess the relationship between all three data types and between methylation and expression, controlling for the effect of genetic distance (Vegan, Oksanen *et al.* 2017).

We also obtained the probability of whether significantly associated SNPs and methylation positions were likely to be overlapping differentially expressed genes (DEGs) by chance using a bootstrap method. We drew a number of random SNPs or methylated positions, with replacement, equal to the number of observed significantly associated SNPs or DMPs, and counted the number of loci that overlapped with DEGs in our previous study. This process was repeated 9999 times each for genetic and methylation data. We derived P-values by counting the number of times a value at least as large as the observed value appeared in the bootstrap resamples and dividing by the number of bootstrap replicates.

### Greenhouse oil response experiment

We assessed the possibility that native *S. alterniflora* populations harbored genetic variation for the response to crude oil via a controlled greenhouse experiment. We collected 10 *S. alterniflora* individuals that had been collected from two oil-naïve sites (“Cabretta” and “Lighthouse”) in the Sapelo Island National Estuarine Research Reserve in Georgia, USA, in May 2010. We collected plants that were spaced ten meters apart, maximizing the chance that individuals were of different genotypes (Foust *et al.*, 2016; Richards, Hamrick, Donovan, & Mauricio, 2004). We grew these individuals in pots under greenhouse conditions for at least three years before beginning our experiments and propagated multiple replicates of each genotype by rhizome cutting. Individual ramets were separated and potted in 4 inch pots in a 50-50 mixture of peat and sand (Cypress Creek, Tampa, USA; Alvarez, 2016).

We distributed three potted replicates of each of the 10 genotypes in each of two plastic containment chambers, for a total of 60 biological samples. One chamber received only uncontaminated fresh water, while the oil treatment chamber received 2.5% oil (sweet Louisiana crude) in 62 liters of water, which we previously determined would induce strong phenotypic response (Alvarez, 2016). Tides were simulated once per day by filling containment chambers with the water or water-oil mixture and allowing the fluid to drain into a catchment. We estimated biomass by tallying the number of living leaves and the number of living ramets when the experiment began, and again seven days after crude oil was added.

### Statistical analysis of greenhouse experiment

We used generalized linear models (R Core Team 2017) with a Poisson error distribution to analyze the above-ground biomass at the end of the experiment (measured as the number of leaves and the number of ramets). Because *S. alterniflora* reproduces clonally, we assumed that biomass would represent a reasonable proxy of fitness in our species (Younginger, Sirová, Cruzan, & Ballhorn, 2017). We also included a covariate for the size of each plant at the start of the experiment, represented by the number of leaves and the number of ramets at the start of the experiment. Each model was written as (Phenotype_Final_ ~ Treatment * Genotype + Phenotype_Initial_), where asterisks represent main effects and all interactions. We did not explicitly test for differences between sites since admixture is high between sites on Sapelo Island and we found no evidence of genetic differentiation (Foust *et al.*, 2016). We assessed significance for main effects and interactions using type III tests. However, to identify individual genotypes responding more strongly than others, we conducted post-hoc pairwise comparisons, correcting for multiple testing using Holm’s correction for multiple testing (emmeans; Lenth, 2018).

## Results

### epiGBS yields informative genetic and methylation loci

The libraries for 48 individuals (Table 1) generated 6,809,826 total raw sequencing reads, of which 3,833,653 (56.3%) could be matched to their original mate strand. *De novo* assembly using the epiGBS pipeline resulted in 36,131 contiguous fragments ranging from 19-202 bp, an average length of 123.92 bp, and a total length of 5,441,437 bp. The size of the *S. alterniflora* genome is estimated to be 2C values = 6x = 4.2 GB (Fortune *et al.*, 2008), and current genomic analyses indicate that repetitive sequences (including transposable elements and tandem repeats) represent about 45% of the total analyzed genomic data set in *S. alterniflora* (Giraud *et al., in prep).* Therefore, we estimate that our epiGBS approach assayed approximately 0.6% of the non-repetitive genome. However, fragments that were >90% similar were merged, and polyploid homeologs may have been concatenated. With BLAST, we found 10,103 fragments mapped to 2,718 transcripts in the *S. alterniflora* transcriptome. We found that 1,571 transcripts (57.8%) contained multiple epiGBS fragments that align to the same place, and 296 (10.9%) contained multiple epiGBS fragments that mapped to different places within the same gene. We suspect that multiple epiGBS fragments map to the same location because some epiGBS fragments represent different homeologs of the same region, which mapped to the same location. Only 1% of reads map to repetitive elements, confirming that *Pst1*-fragmented libraries were biased away from highly methylated, repetitive regions (van Gurp *et al.*, 2016). The bisulfite non-conversion rate was calculated to be 0.36% of cytosines per position, and was estimated from lambda phage spike-in (van Gurp *et al.*, 2016). Although we found substantial variation in average sequencing depth among samples, we found no obvious non-random bias in sampling depth across samples (Figure S1). However, during filtering, we removed ten samples due to stochastic under-sequencing, leaving 38 samples for population analyses (Table 1, Figure S2).

### Genetic differentiation

Our initial sequencing run yielded 171,205 SNPs across all individuals. After filtering to common loci, removing invariant sites, and imputing missing data (Figure S2), we obtained 63,796 SNP loci. Of these, 5,753 SNPs occurred in transcripts. As in our AFLP study, we found significant genetic differentiation that was correlated to oil exposure: oil exposure explained 23.4% of the variance in DNA sequence (P<0.001, Figure 1A, B, Table 2), providing evidence that selection may have acted on these populations. Pairwise F_ST_ calculations showed that all sites were significantly different from each other (Table 3), with no evidence of isolation by distance (P>0.05 for latitude, longitude, and interaction). We found 1,631 SNPs that were significantly associated with oil exposure (defined by a locus score >3 standard deviation units away from the mean locus score; Forester *et al.*, 2018; Figure 1C), including 169 that overlapped with the *S. alterniflora* transcriptome. Of these loci, 41 were annotated using information from *O. sativa,* and contained a number of putative regulators of gene expression. Among significant loci, 1,324 differed in major allele frequency between exposed and unexposed populations by greater than 5%, and 334 by greater than 20% (Figure 1C). Significantly differentiated loci appeared no less likely to increase or decrease in major allele frequency based on exposure (809 increasing vs 822 decreasing in frequency). We tested for homogeneity of group dispersion, and found no evidence of change in variance in oil-exposed populations (P=0.512).

**Table 2.** Association between oil exposure, genetic distance, and methylation distance across tests. Test statistics and significance determined through RDA. *** indicates significance at *P* < 0.001.

|  | df | F |  | Variance Explained |
| --- | --- | --- | --- | --- |
| Genetic | 1 | 2.183 | *** | 0.234 |
| Methylation |  |  |  |  |
| Without control for genetic var. | 1 | 1.96 | *** | 0.167 |
| With control for genetic var. | 1 | 1.199 |  | 0.099 |

**Table 3.**
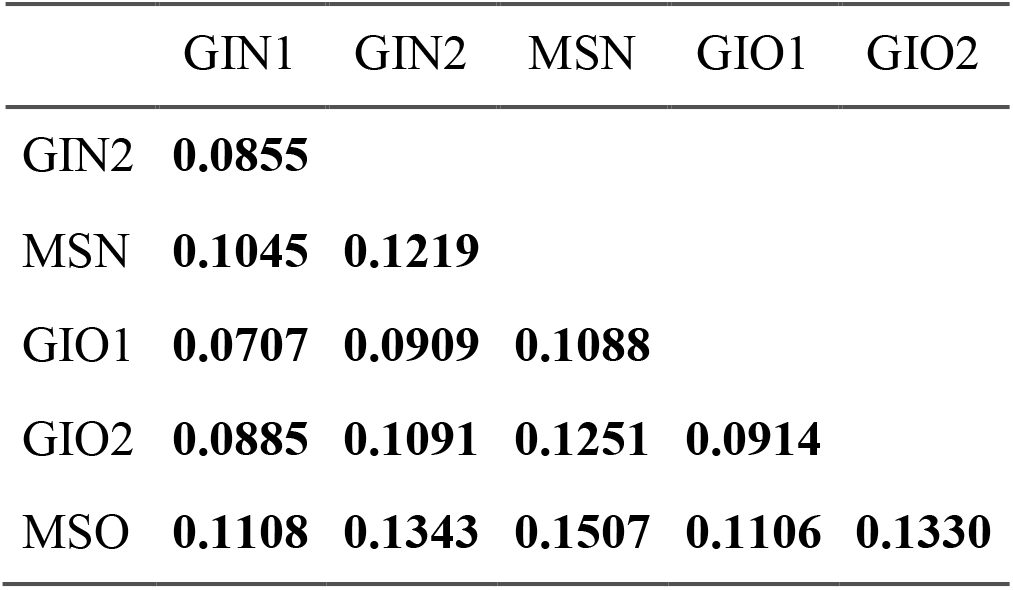
Pairwise F_ST_ among three oil contaminated and three uncontaminated sites. Bold (i.e. all entries) indicates significance at *P* < 0.001.

### DNA methylation differentiation

Our bisulfite sequencing yielded 1,402,083 cytosines that were polymorphic for methylation across our samples before filtering. After filtering our data to common loci as described above, we analyzed 92,999 polymorphic methylated loci, 25,381 of which occurred in the CG context, 24,298 in the CHG context, and 43,030 in the CHH context (Figure 2C). An additional 290 had variable context because they co-occurred with a SNP. These proportions of polymorphic methylation loci did not change significantly due to oil exposure. Methylation calls were collapsed for symmetric CG and CHG loci across “watson” and “crick” strands so that methylation on either one or both strands was considered as a single locus. Although DNA methylation was strongly correlated with oil exposure (Table 2, P<0.001) when controlling only for latent factors, this differentiation was not significant after controlling for genetic population structure with principal components of genetic data (Table 2, P>0.1). In the latter model, oil explained 10% of the variation in methylation.

We found 240 DMPs that differed significantly between exposure types (Q<0.05, Figure 2C; Table S1). The number of observed DMPs in the CG context (125 loci) was significantly overrepresented relative to their occurrence in our data (P<0.001). We also found DMPs in CHG (57 loci), and CHH (58 loci) context, which was underrepresented among DMPs relative to their prevalence in all contigs (P<0.001). Among the significant loci, most had negligible differences in the magnitude of methylation frequency changes (average 1.4% change between exposed and unexposed populations). Only 29 experienced a change in magnitude of methylation greater than 5%, and only 7 loci showed a change of greater than 20%. Additionally, 19 DMPs were located within a fragment that mapped to the *S. alterniflora* transcriptome, and 49 DMPs occurred in the same fragment as a significantly differentiated SNP. However, only 4 of those SNPs altered the trinucleotide context of DNA methylation.

### Correlations with gene expression

We found no significant relationship between genetic distance and gene expression distance (Mantel’s R= 0.050, P=0.32), between patterns of methylation variation and genome wide gene expression (Mantel’s R=0.051, P=0.29), or between methylation and genome wide expression when controlling for genetic variation (Mantel’s R=0.014, P= 0.41). Only 14 SNPs that were significantly associated with oil exposure overlapped with DEGs correlated with exposure to the *DWH* oil spill (Alvarez *et al.*, 2018). However, our bootstrap test showed that this overlap could occur by chance (P>0.79). Therefore, our data suggests that if these SNPs are under selection, they are not necessarily regulating differential expression resulting from coding changes in those genes. In addition, although 19 DMPs overlapped coding regions, only 3 of the DMPs corresponded to a DEG (Table S1), and our bootstrap test suggests that this was also likely to occur by chance (P>0.5). However, our data is limited to address the association between DNA methylation and differential expression of specific genes.

### Genotypes in common garden differ in their response to crude oil

We found a significant effect of oil exposure on both the number of leaves (F= 13.09, P < 0. 001) and the number of ramets (F = 28.75, P < 0.001) at the end of the controlled greenhouse experiment. Type III tests showed significant genotype-by-treatment interactions for the number of leaves, but not ramets, at the end of the experiment, suggesting that individual genotypes vary in their response to crude oil exposure. Post-hoc comparisons identified two genotypes (C and G; FDR<0.05, Figure 3; Table S2) that lost a significantly greater number of leaves over the course of the experiment relative to other genotypes, further suggesting the presence of standing variation among individuals for the response to crude oil exposure.

**Figure 3.**
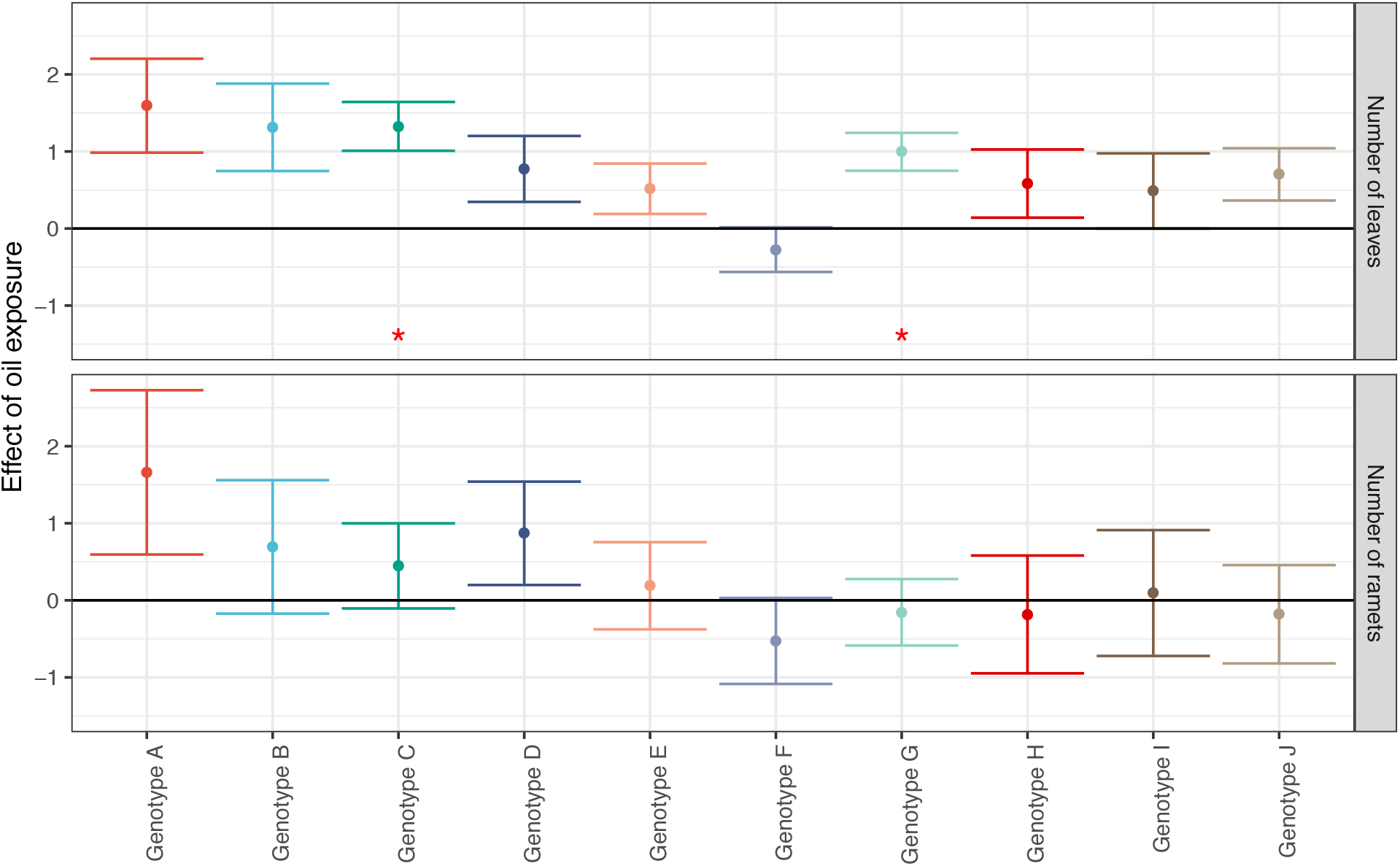
Variation in effect size estimates of the effect of crude oil exposure in individual genotypes. Estimates were based on estimated marginal means in our greenhouse experiment. Asterisks represent significance in post-hoc comparisons.

## Discussion

*Spartina alterniflora* displays high levels of genetic and DNA methylation variation across environmental conditions in its native range (Foust *et al.*, 2016; Hughes & Lotterhos, 2014; Richards *et al.*, 2004; Robertson *et al.*, 2017), potentially providing substrate for both genetic and epigenetic response to pollution. We previously found that genetic structure and expression of 3,334 genes were correlated to exposure to the *DWH* oil spill, but genome-wide methylation variation was not (Alvarez *et al.*, 2018; Robertson *et al.*, 2017). Higher resolution epiGBS suggests that both genetic sequence and DNA methylation are correlated with crude oil exposure in *S. alterniflora,* but that differentiation in DNA methylation is primarily explained by differences in allele frequencies. Additionally, our greenhouse experiment shows phenotypic plasticity and genotypic variation in crude oil response, as measured by differential reduction in biomass between exposed and unexposed samples. These findings are consistent with genotype-specific mortality, and suggest that the *DWH* oil spill may have been a selective event in *S. alterniflora* populations.

### Genetic and epigenetic response to the DWH

We found significant genetic differentiation between oil-exposed and unexposed sites, which may reflect either stochastic mortality in oil-exposed areas from a severe bottleneck, or a signature of selection for oil tolerance in affected populations. *Spartina alterniflora* displays high phenotypic plasticity, and populations have persisted after exposure to the *DWH* oil spill, even after extensive aboveground dieback (Lin & Mendelssohn, 2012; Lin *et al.*, 2016; Silliman *et al.*, 2012). However, our studies and previous accounts of initial losses in live aboveground and belowground biomass (Lin *et al.*, 2016) suggest that some *S. alterniflora* genotypes were more susceptible than others to crude oil stress, and either had not regrown at the time of sampling or experienced mortality as a result of oil exposure. Although we found no evidence for a reduction in genetic variation, which may have further indicated selection for tolerant genotypes, the high ploidy of *S. alterniflora* makes accurate quantification of total genetic variation challenging. Further investigations are required to confirm the magnitude of selection, whether mortality varied by genotype, and if there was a reduction in genetic variation among oil-exposed populations.

DNA methylation differences may reflect either the downstream effects of genetic variants, an induced response to environment, or both (Meng *et al.*, 2016). For example, in another study of *S. alterniflora* populations, patterns of DNA methylation were more strongly correlated than genetic structure with microhabitat, and correlation of DNA methylation to environment was independent of population structure (Foust *et al.*, 2016). In this study, we found a multi-locus signature of methylation differentiation (17% of the variation explained by oil exposure) between oil-affected and unaffected sites before controlling for population structure. However, we found no association between methylation and crude oil contamination after controlling for genetic variation and latent effects, suggesting DNA methylation is controlled by genetic variation.

The observed variation in DNA methylation may be controlled by genetic variation via either a change in the nucleotide context, the presence or absence of particular alleles in *cis,* or variation in upstream regulatory elements. Allelic variation that changes trinucleotide context can alter or eliminate the ability of a methyltransferase to deposit methylation at that site. However, in our data, we did not find an enrichment of SNPs that affected trinucleotide context of DMPs. Concurrently, we did not detect an enrichment of oil-associated SNPs in DEGs, which we would expect if changes in the coding regions of those genes explain the observed gene expression variation in oil-exposed individuals. However, our ability to assess the relationships between SNPs, SMPs and DEGs was limited by the distribution of our RRBS fragments. Further, changes in allele frequencies, due to either selection or drift, may have generated divergence in the regulatory machinery maintaining DNA methylation and gene expression profiles among exposed and unexposed populations.

Although we cannot disentangle whether differential expression causes alternative methylation patterns or vice versa, we previously identified a DEG that was homologous to the histone methyltransferase SUVH5, which may modulate fitness effects during oil exposure (Alvarez *et al.*, 2018). Histone methylation is linked to DNA methylation through the regulation of CHROMOMETHYLASE3 (CMT3) activity (Stroud, Greenberg, Feng, Bernatavichute, & Jacobsen, 2013). Given our previous results and those from the present study, we hypothesized that the differential expression of SUVH5 in response to crude oil exposure would result in differences in DNA methylation. These differences, in turn, may be maintained via genetic variation between exposed and unexposed populations either in the SUVH5 homolog itself, or more broadly within the CMT3-mediated pathway. However, targeted resequencing and further functional validation in the populations of interest will be required to confirm this hypothesis.

### Reduced representation sequencing compared to AFLP

As the field of ecological genomics matures, there is a pressing need to develop robust assays and statistically sound measures of regulatory variation. Reduced representation methylation sequencing techniques are attractive for ecological epigenomics because they can be used to infer genome wide patterns of both genetic and methylation variation without a high-quality reference genome (Paun, Verhoeven, & Richards, 2019; Richards *et al.*, 2017; Robertson & Richards, 2015; van Moorsel *et al.* 2019). However, they still have serious limitations particularly for species that do not yet have a fully sequenced reference genome (Paun *et al.*, 2019). Furthermore, it is important to note that the limited number of loci surveyed may have led to a biased subsampling of the genome. In turn, this can lead to a poor estimation of the “neutral” level of divergence in the genome, and therefore a biased interpretation of divergence between these populations (Lowry *et al*., 2017).

When comparing epiGBS to MS-AFLP, we expected that the substantial increase in markers (92,999 compared to 39 polymorphic methylation loci, respectively) would lend greater resolution to detect patterns of DNA methylation variation. Our epiGBS survey detected significant differentiation in both genetic variation and DNA methylation that was correlated to oil exposure, suggesting that epiGBS provides increased resolution over MS-AFLP to detect genome-wide methylation differences. However, despite the much larger data set generated by epiGBS, we only found 240 differentially methylated positions. Although it would be valuable to identify associations between gene expression and nearby DNA methylation variation, the minimal overlap between our RRBS fragments and DEGs hindered our ability to associate methylation and gene expression variation. This is due to the small fraction of the genome that is assayed, substantial variation in methylation, and that we were unable to identify fragments that overlapped promoter regions without a reference genome.

Future RRBS studies will benefit from optimizing protocols that enrich for specific portions of the genome (e.g. Heer & Ullrich *et al.*, 2018), but generating a draft reference genome will be imperative to allow for better exploitation of RRBS data and ascertain gene function (Paun *et al.*, 2019). While sequencing-based techniques provide the potential to identify functional genomic regions, correct annotations rely on genomic resources in a relevant reference. In polyploid species like *S. alterniflora,* the number of duplicated genes and the potential for neofunctionalization among them creates additional uncertainty for correctly assigning annotations (Primmer, Papkostas, Leder, Davis & Ragan, 2013). *Spartina alterniflora* has various levels of duplicated gene retention, small RNA variation (including miRNAs and SiRNAs) and homeologous expression (Ainouche, Baumel, Salmon, & Yannic 2003; Boutte *et al.*, 2016; Cavé-Radet, Giraud, Lima, El Amrani, Ainouche, & Salmon, 2019; Ferreira de Carvalho, 2013, 2017; Fortune *et al.*, 2007), which may result in more opportunities for gene diversification and subfunctionalization (Chen *et al.*, 2015; Salmon & Ainouche, 2015; Shimizu-Inatsugi *et al.*, 2017). Therefore, while studies with RRBS techniques in non-model plants offer increased power to detect broad, genome-wide patterns of variation that may be correlated to ecology, they are still limited for the detection of specific gene function. Improving genomics resources in a variety of organisms is an essential next step for understanding the molecular level basis of ecological interactions.

## Acknowledgments

We thank Steve Pennings, Brittney DeLoach McCall, Aaron Schrey, Christina Moss, and Ashley Shayter for access to field sites and assistance with plant sampling. We thank Acer Van Wallendael, Kieran Samuk, and Kate Ostevik for constructive feedback. This work was supported by funding from the National Science Foundation (U.S.A.) DEB-1419960 and IOS-1556820 (to CLR) and through the Global Invasions Network Research Exchange (Grant No. 0541673 for MR), the Franco-American Fulbright Commission (to CLR), and the Netherlands Organisation for Scientific Research (NWO-ALW No. 820.01.025 to KJFV).

## Author Contributions

CLR & KJFV conceived the study. CLR, KJFV, MA, MR, MLA and AS designed the experiments and analyses. MA, MR, CAMW and TVG did the laboratory work. MA, MR, and TVG analyzed the epiGBS data. CLR, MA, DG, and AS analyzed the transcriptome and gene expression data. CLR, MA and MR wrote the first draft of the manuscript. All co-authors provided input and revisions to the manuscript.

## Data accessibility

Raw data files are available on Dryad at XXX. Supplementary tables and figures can be found in the electronic supplementary material, and is available on github.com/alvarezmf/DWH_epigbs along with processed data and R scripts.

## Supporting Information

Table S1 SNPs and methylation loci significantly associated with oil exposure.

Table S2 Post-hoc tests for the effect of crude oil exposure in individual genotypes. Estimates were based on estimated marginal means in our greenhouse experiment.

Figure S1 A) Principal components analysis on sampling depth per SNP allele. B) Percent variance explained by each principal component.

Figure S2 Upper half: percentage of present and imputed data per sample after filtering for A) SNP and B) methylation loci. Lower half: percentage of present and imputed data per locus after filtering for C) SNP and D) methylation loci. In C and D, only the first 5,000 loci are shown for clarity. Dotted lines represent missing data cut-off values for removal (80% missing for individuals in A & B, 50% for loci in C & D).

