## Supplementary material for "Reduced representation characterization of genetic and epigenetic differentiation to oil pollution in the foundation plant *Spartina alterniflora*": Table S2

TableS2

| Dependent | contrast | Genotype | estimate | SE | df | z.ratio | p.value | holm |
|---|---|---|---|---|---|---|---|---|
| **Number of leaves** | Control - OilTreatment | Genotype A | 1.5941718323692 | 0.609475297046669 | Inf | 2.616 | 0.009 | 0.16 |
| **Number of leaves** | Control - OilTreatment | Genotype B | 1.31363805651122 | 0.567039246638814 | Inf | 2.317 | 0.021 | 0.349 |
| **Number of leaves** | Control - OilTreatment | Genotype C | 1.32615647584022 | 0.316743999969456 | Inf | 4.187 | 0 | 0.001 |
| **Number of leaves** | Control - OilTreatment | Genotype D | 0.77359848768348 | 0.428137556311393 | Inf | 1.807 | 0.071 | 1 |
| **Number of leaves** | Control - OilTreatment | Genotype E | 0.516011583301125 | 0.327234450244056 | Inf | 1.577 | 0.115 | 1 |
| **Number of leaves** | Control - OilTreatment | Genotype F | -0.273884060865239 | 0.289994848577945 | Inf | -0.944 | 0.345 | 1 |
| **Number of leaves** | Control - OilTreatment | Genotype G | 0.995385711552419 | 0.246071755151921 | Inf | 4.045 | 0 | 0.001 |
| **Number of leaves** | Control - OilTreatment | Genotype H | 0.583362844287245 | 0.442402075933546 | Inf | 1.319 | 0.187 | 1 |
| **Number of leaves** | Control - OilTreatment | Genotype I | 0.487010820767995 | 0.488302862009079 | Inf | 0.997 | 0.319 | 1 |
| **Number of leaves** | Control - OilTreatment | Genotype J | 0.701997378865003 | 0.339318203390244 | Inf | 2.069 | 0.039 | 0.617 |
| **Number of ramets** | Control - OilTreatment | Genotype A | 1.660563229318 | 1.06602018569789 | Inf | 1.558 | 0.119 | 1 |
| **Number of ramets** | Control - OilTreatment | Genotype B | 0.693147180559943 | 0.866025388389517 | Inf | 0.8 | 0.423 | 1 |
| **Number of ramets** | Control - OilTreatment | Genotype C | 0.446824315789697 | 0.552014816134584 | Inf | 0.809 | 0.418 | 1 |
| **Number of ramets** | Control - OilTreatment | Genotype D | 0.869615887892201 | 0.670570795272524 | Inf | 1.297 | 0.195 | 1 |
| **Number of ramets** | Control - OilTreatment | Genotype E | 0.189261300898759 | 0.56595873458857 | Inf | 0.334 | 0.738 | 1 |
| **Number of ramets** | Control - OilTreatment | Genotype F | -0.527024505714794 | 0.558119159986764 | Inf | -0.944 | 0.345 | 1 |
| **Number of ramets** | Control - OilTreatment | Genotype G | -0.155690653406628 | 0.431530487742454 | Inf | -0.361 | 0.718 | 1 |
| **Number of ramets** | Control - OilTreatment | Genotype H | -0.182970883422871 | 0.764380624490837 | Inf | -0.239 | 0.811 | 1 |
| **Number of ramets** | Control - OilTreatment | Genotype I | 0.0947785987780016 | 0.816884488038075 | Inf | 0.116 | 0.908 | 1 |
| **Number of ramets** | Control - OilTreatment | Genotype J | -0.18059589873312 | 0.638310179024628 | Inf | -0.283 | 0.777 | 1 |
