## Supplementary material for "Reduced representation characterization of genetic and epigenetic differentiation to oil pollution in the foundation plant *Spartina alterniflora*": Table S1

TableS1

| Contig | Position | Type | Microarray ID | Transcriptome ID | Oryza sativa ID |
|---|---|---|---|---|---|
| **C:11159** | 82 | SNP | | Salt_2_contig10017 | LOC_Os09g38620 |
| **C:2737** | 6 | SNP | | Salt_2_contig10546 | LOC_Os02g31220 |
| **C:1215** | 80 | SNP | | Salt_2_contig10909 | |
| **C:433** | 72 | SNP | | Salt_2_contig10909 | |
| **C:433** | 62 | SNP | | Salt_2_contig10909 | |
| **C:433** | 107 | SNP | | Salt_2_contig10909 | |
| **C:433** | 24 | CHG | | Salt_2_contig10909 | |
| **C:433** | 88 | CG | | Salt_2_contig10909 | |
| **C:4056** | 185 | SNP | | Salt_2_contig11658 | |
| **C:4056** | 129 | SNP | | Salt_2_contig11658 | |
| **C:30772** | 28 | SNP | | Salt_2_contig12104 | |
| **C:2865** | 6 | SNP | | Salt_2_contig12302 | |
| **C:2865** | 109 | SNP | | Salt_2_contig12302 | |
| **C:2865** | 86 | SNP | | Salt_2_contig12302 | |
| **C:1253** | 9 | SNP | | Salt_2_contig13172 | |
| **C:355** | 74 | SNP | | Salt_2_contig13182 | |
| **C:16** | 133 | CHH | | Salt_2_contig13533 | LOC_Os09g24710 |
| **C:1654** | 197 | SNP | | Salt_2_contig13814 | |
| **C:1654** | 133 | SNP | | Salt_2_contig13814 | |
| **C:10606** | 47 | SNP | | Salt_2_contig13834 | |
| **C:5064** | 96 | CG | | Salt_2_contig13839 | |
| **C:703** | 170 | SNP | S_5sp_contig28578 | Salt_2_contig142 | |
| **C:12768** | 26 | SNP | | Salt_2_contig16439 | |
| **C:484** | 141 | SNP | | Salt_2_contig166 | LOC_Os09g07500 |
| **C:484** | 144 | SNP | | Salt_2_contig166 | LOC_Os09g07500 |
| **C:4904** | 23 | SNP | | Salt_2_contig16786 | LOC_Os02g49332 |
| **C:744** | 167 | SNP | | Salt_2_contig17047 | |
| **C:736** | 195 | SNP | | Salt_2_contig17352 | |
| **C:1805** | 119 | SNP | | Salt_2_contig18090 | LOC_Os01g58790 |

| | | | | | |
|---|---|---|---|---|---|
| **C:1805** | 196 | SNP | | Salt_2_contig18090 | LOC_Os01g58790 |
| **C:11665** | 12 | SNP | | Salt_2_contig1822 | |
| **C:11268** | 162 | SNP | | Salt_2_contig18793 | |
| **C:305** | 175 | SNP | | Salt_2_contig189 | |
| **C:305** | 65 | SNP | | Salt_2_contig189 | |
| **C:305** | 169 | SNP | | Salt_2_contig189 | |
| **C:305** | 160 | SNP | | Salt_2_contig189 | |
| **C:27859** | 104 | SNP | | Salt_2_contig18977 | |
| **C:27859** | 104 | SNP | | Salt_2_contig18977 | |
| **C:27859** | 104 | SNP | | Salt_2_contig18977 | LOC_Os01g57968 |
| **C:27859** | 104 | SNP | | Salt_2_contig18977 | |
| **C:27859** | 104 | SNP | | Salt_2_contig18977 | |
| **C:27859** | 104 | SNP | | Salt_2_contig18977 | |
| **C:27859** | 104 | SNP | | Salt_2_contig18977 | |
| **C:27859** | 104 | SNP | | Salt_2_contig18977 | |
| **C:27859** | 104 | SNP | | Salt_2_contig18977 | |
| **C:27859** | 104 | SNP | | Salt_2_contig18977 | |
| **C:417** | 38 | SNP | | Salt_2_contig20036 | |
| **C:1353** | 112 | SNP | | Salt_2_contig20223 | |
| **C:895** | 20 | SNP | | Salt_2_contig20223 | |
| **C:3765** | 84 | SNP | | Salt_2_contig21188 | |
| **C:4428** | 75 | SNP | | Salt_2_contig21253 | LOC_Os04g42470 |
| **C:27609** | 101 | SNP | | Salt_2_contig22140 | |
| **C:2434** | 166 | SNP | | Salt_2_contig22143 | |
| **C:6045** | 52 | SNP | | Salt_2_contig22352 | |
| **C:1525** | 40 | SNP | | Salt_2_contig22544 | |
| **C:3405** | 135 | SNP | | Salt_2_contig23828 | |
| **C:2906** | 159 | SNP | | Salt_2_contig24963 | |
| **C:2906** | 118 | SNP | | Salt_2_contig24963 | |
| **C:1376** | 47 | SNP | | Salt_2_contig25385 | LOC_Os03g22890 |
| **C:1376** | 111 | SNP | | Salt_2_contig25385 | LOC_Os03g22890 |
| **C:1376** | 95 | SNP | | Salt_2_contig25385 | LOC_Os03g22890 |

| | | | | | |
|---|---|---|---|---|---|
| **C:1376** | 156 | SNP | | Salt_2_contig25385 | LOC_Os03g22890 |
| **C:413** | 74 | SNP | | Salt_2_contig2541 | |
| **C:230** | 134 | SNP | S_5sp_contig44897 | Salt_2_contig2616 | |
| **C:466** | 23 | CHG | | Salt_2_contig26222 | |
| **C:466** | 162 | SNP | | Salt_2_contig26222 | |
| **C:2660** | 116 | SNP | | Salt_2_contig26796 | LOC_Os03g08870 |
| **C:14635** | 113 | SNP | | Salt_2_contig27228 | |
| **C:11363** | 133 | SNP | | Salt_2_contig27262 | LOC_Os01g55880 |
| **C:11363** | 133 | SNP | | Salt_2_contig27262 | LOC_Os08g07540 |
| **C:11363** | 133 | SNP | | Salt_2_contig27262 | |
| **C:1978** | 180 | SNP | | Salt_2_contig27403 | LOC_Os06g05660 |
| **C:1978** | 195 | SNP | | Salt_2_contig27403 | LOC_Os06g05660 |
| **C:10215** | 6 | SNP | | Salt_2_contig27403 | LOC_Os06g05660 |
| **C:10215** | 115 | SNP | | Salt_2_contig27403 | LOC_Os06g05660 |
| **C:5066** | 169 | SNP | S_alt_contig07871 | Salt_2_contig28368 | |
| **C:270** | 180 | SNP | | Salt_2_contig2897 | LOC_Os03g29170 |
| **C:270** | 90 | SNP | | Salt_2_contig2897 | LOC_Os03g29170 |
| **C:510** | 149 | SNP | | Salt_2_contig29452 | LOC_Os04g45470 |
| **C:1045** | 26 | CG | | Salt_2_contig29548 | |
| **C:31248** | 6 | SNP | | Salt_2_contig29810 | LOC_Os07g44940 |
| **C:2419** | 190 | SNP | | Salt_2_contig29818 | |
| **C:2419** | 118 | SNP | | Salt_2_contig29818 | |
| **C:4174** | 119 | SNP | S_5sp_contig45052 | Salt_2_contig30241 | |
| **C:4784** | 8 | SNP | | Salt_2_contig30539 | |
| **C:4784** | 62 | SNP | | Salt_2_contig30539 | |
| **C:11832** | 60 | SNP | | Salt_2_contig30871 | |
| **C:34487** | 13 | SNP | | Salt_2_contig31153 | |
| **C:4671** | 174 | SNP | S_alt_contig12637 | Salt_2_contig31184 | |
| **C:10909** | 101 | SNP | | Salt_2_contig32043 | |
| **C:10909** | 100 | SNP | | Salt_2_contig32043 | |
| **C:1992** | 15 | SNP | | Salt_2_contig32210 | |
| **C:35001** | 113 | SNP | | Salt_2_contig32343 | |

| | | | | | |
|---|---|---|---|---|---|
| **C:1215** | 80 | SNP | | Salt_2_contig32643 | |
| **C:35268** | 94 | SNP | | Salt_2_contig32813 | |
| **C:11155** | 104 | SNP | S_5sp_contig41202 | Salt_2_contig33117 | LOC_Os03g03650 |
| **C:11156** | 8 | SNP | S_5sp_contig41202 | Salt_2_contig33117 | LOC_Os03g03650 |
| **C:30048** | 36 | SNP | | Salt_2_contig33536 | LOC_Os03g44780 |
| **C:28767** | 73 | SNP | | Salt_2_contig34798 | |
| **C:28767** | 62 | SNP | | Salt_2_contig34798 | |
| **C:8120** | 57 | SNP | | Salt_2_contig34823 | |
| **C:1304** | 160 | SNP | | Salt_2_contig35877 | LOC_Os01g73790 |
| **C:9841** | 91 | SNP | | Salt_2_contig35980 | |
| **C:9855** | 98 | SNP | | Salt_2_contig35980 | |
| **C:29258** | 12 | SNP | | Salt_2_contig36339 | |
| **C:5057** | 70 | SNP | | Salt_2_contig3646 | |
| **C:10239** | 63 | SNP | | Salt_2_contig36647 | LOC_Os02g42890 |
| **C:10239** | 60 | SNP | | Salt_2_contig36647 | LOC_Os02g42890 |
| **C:10239** | 43 | SNP | | Salt_2_contig36647 | LOC_Os02g42890 |
| **C:2446** | 15 | SNP | | Salt_2_contig3707 | |
| **C:7392** | 85 | SNP | | Salt_2_contig3714 | |
| **C:1121** | 189 | SNP | S_alt_contig04187 | Salt_2_contig37963 | LOC_Os06g10760 |
| **C:1557** | 63 | SNP | | Salt_2_contig38269 | |
| **C:3644** | 141 | SNP | | Salt_2_contig38713 | |
| **C:7636** | 9 | SNP | | Salt_2_contig39997 | LOC_Os12g13150 |
| **C:11** | 61 | CHH | S_5sp_contig38692 | Salt_2_contig40062 | LOC_Os02g12350 |
| **C:11** | 61 | CHH | S_5sp_contig38692 | Salt_2_contig40062 | LOC_Os06g38470 |
| **C:11** | 26 | SNP | S_5sp_contig38692 | Salt_2_contig40062 | LOC_Os02g12350 |
| **C:11** | 26 | SNP | S_5sp_contig38692 | Salt_2_contig40062 | LOC_Os06g38470 |
| **C:14044** | 15 | SNP | S_alt_contig06820_F | Salt_2_contig40110 | |
| **C:11533** | 103 | SNP | | Salt_2_contig40120 | |
| **C:4617** | 82 | SNP | | Salt_2_contig40149 | |
| **C:4617** | 108 | SNP | | Salt_2_contig40149 | |
| **C:2052** | 111 | SNP | | Salt_2_contig40154 | |
| **C:2052** | 197 | SNP | | Salt_2_contig40154 | |

| | | | | | |
|---|---|---|---|---|---|
| **C:35830** | 17 | SNP | | Salt_2_contig40261 | |
| **C:633** | 67 | SNP | | Salt_2_contig40278 | LOC_Os02g47970 |
| **C:9302** | 126 | SNP | | Salt_2_contig40290 | LOC_Os01g15630 |
| **C:373** | 51 | CHG | | Salt_2_contig40331 | LOC_Os06g04270 |
| **C:1024** | 48 | SNP | | Salt_2_contig40421 | LOC_Os02g54500 |
| **C:4808** | 132 | SNP | | Salt_2_contig40711 | LOC_Os02g40430 |
| **C:4808** | 132 | SNP | | Salt_2_contig40711 | LOC_Os04g42840 |
| **C:645** | 51 | SNP | | Salt_2_contig40729 | |
| **C:645** | 45 | SNP | | Salt_2_contig40729 | |
| **C:3703** | 6 | SNP | | Salt_2_contig41307 | LOC_Os04g52900 |
| **C:3703** | 43 | SNP | | Salt_2_contig41307 | LOC_Os04g52900 |
| **C:11137** | 140 | SNP | | Salt_2_contig41353 | LOC_Os05g25060 |
| **C:1661** | 192 | CHG | | Salt_2_contig41409 | |
| **C:14018** | 106 | SNP | | Salt_2_contig41431 | LOC_Os08g43130 |
| **C:14018** | 129 | SNP | | Salt_2_contig41431 | LOC_Os08g43130 |
| **C:14018** | 19 | SNP | | Salt_2_contig41431 | LOC_Os08g43130 |
| **C:14018** | 84 | SNP | | Salt_2_contig41431 | LOC_Os08g43130 |
| **C:14018** | 116 | SNP | | Salt_2_contig41431 | LOC_Os08g43130 |
| **C:1175** | 126 | SNP | | Salt_2_contig41459 | |
| **C:1905** | 47 | SNP | | Salt_2_contig41630 | LOC_Os03g51020 |
| **C:1905** | 140 | SNP | | Salt_2_contig41630 | LOC_Os03g51020 |
| **C:1905** | 131 | SNP | | Salt_2_contig41630 | LOC_Os03g51020 |
| **C:1905** | 120 | SNP | | Salt_2_contig41630 | LOC_Os03g51020 |
| **C:1622** | 122 | SNP | S_5sp_contig10902 | Salt_2_contig41793 | |
| **C:47** | 135 | SNP | | Salt_2_contig42072 | LOC_Os02g19220 |
| **C:47** | 32 | CHG | | Salt_2_contig42072 | LOC_Os02g19220 |
| **C:11839** | 58 | SNP | | Salt_2_contig42190 | LOC_Os03g11340 |
| **C:35701** | 61 | SNP | | Salt_2_contig42391 | |
| **C:30251** | 132 | CG | | Salt_2_contig42856 | LOC_Os04g41100 |
| **C:30251** | 132 | CG | | Salt_2_contig42856 | LOC_Os02g39010 |
| **C:6795** | 102 | SNP | | Salt_2_contig43059 | |
| **C:1252** | 31 | SNP | | Salt_2_contig43111 | LOC_Os04g36660 |

| | | | | | |
|---|---|---|---|---|---|
| **C:1226** | 139 | SNP | | Salt_2_contig43158 | |
| **C:265** | 45 | SNP | S_5sp_contig39260 | Salt_2_contig43244 | |
| **C:14319** | 112 | SNP | | Salt_2_contig4326 | |
| **C:106** | 12 | SNP | | Salt_2_contig43323 | LOC_Os11g03160 |
| **C:106** | 12 | SNP | | Salt_2_contig43323 | LOC_Os12g02910 |
| **C:106** | 76 | SNP | | Salt_2_contig43323 | LOC_Os11g03160 |
| **C:106** | 76 | SNP | | Salt_2_contig43323 | LOC_Os12g02910 |
| **C:106** | 87 | SNP | | Salt_2_contig43323 | LOC_Os11g03160 |
| **C:106** | 87 | SNP | | Salt_2_contig43323 | LOC_Os12g02910 |
| **C:714** | 137 | SNP | S_5sp_contig33654 | Salt_2_contig43395 | |
| **C:6718** | 150 | SNP | | Salt_2_contig43636 | |
| **C:6718** | 44 | SNP | | Salt_2_contig43636 | |
| **C:6101** | 109 | SNP | | Salt_2_contig43650 | LOC_Os01g67210 |
| **C:1128** | 159 | SNP | | Salt_2_contig43700 | |
| **C:14697** | 40 | CG | S_5sp_contig32950_ | Salt_2_contig43710 | |
| **C:434** | 131 | SNP | | Salt_2_contig43742 | |
| **C:11486** | 65 | SNP | | Salt_2_contig43832 | |
| **C:11486** | 64 | CG | | Salt_2_contig43832 | |
| **C:1806** | 41 | SNP | | Salt_2_contig43843 | |
| **C:704** | 155 | SNP | | Salt_2_contig43962 | LOC_Os03g42770 |
| **C:704** | 154 | SNP | | Salt_2_contig43962 | LOC_Os03g42770 |
| **C:13895** | 70 | SNP | | Salt_2_contig43962 | LOC_Os03g42770 |
| **C:6668** | 153 | SNP | | Salt_2_contig43974 | |
| **C:1022** | 125 | SNP | | Salt_2_contig4520 | |
| **C:703** | 170 | SNP | S_5sp_contig28578 | Salt_2_contig459 | |
| **C:40** | 20 | SNP | | Salt_2_contig4604 | |
| **C:40** | 159 | SNP | | Salt_2_contig4604 | |
| **C:40** | 81 | SNP | | Salt_2_contig4604 | |
| **C:33810** | 48 | SNP | | Salt_2_contig4907 | LOC_Os02g39890 |
| **C:737** | 61 | SNP | | Salt_2_contig5269 | |
| **C:3580** | 178 | SNP | | Salt_2_contig5419 | |
| **C:11244** | 57 | SNP | S_5sp_contig33054 | Salt_2_contig5455 | |

| | | | | | |
|---|---|---|---|---|---|
| **C:11244** | 56 | SNP | S_5sp_contig33054 | Salt_2_contig5455 | |
| **C:14381** | 34 | SNP | | Salt_2_contig5473 | |
| **C:2695** | 136 | SNP | | Salt_2_contig5936 | |
| **C:11704** | 100 | SNP | | Salt_2_contig5964 | |
| **C:11704** | 97 | SNP | | Salt_2_contig5964 | |
| **C:11704** | 85 | SNP | | Salt_2_contig5964 | |
| **C:6762** | 28 | SNP | | Salt_2_contig6546 | LOC_Os01g07420 |
| **C:6762** | 28 | SNP | | Salt_2_contig6546 | |
| **C:6762** | 28 | SNP | | Salt_2_contig6546 | |
| **C:6762** | 28 | SNP | | Salt_2_contig6546 | |
| **C:6762** | 28 | SNP | | Salt_2_contig6546 | |
| **C:6762** | 28 | SNP | | Salt_2_contig6546 | |
| **C:125** | 156 | CG | S_alt_contig05819 | Salt_2_contig7784 | LOC_Os01g51840 |
| **C:30308** | 78 | CG | | Salt_2_contig793 | |
| **C:30308** | 62 | CG | | Salt_2_contig793 | |
| **C:30308** | 54 | CG | | Salt_2_contig793 | |
| **C:30308** | 114 | CG | | Salt_2_contig793 | |
| **C:705** | 92 | SNP | | Salt_2_contig8321 | |
| **C:705** | 128 | SNP | | Salt_2_contig8321 | |
| **C:705** | 54 | SNP | | Salt_2_contig8321 | |
| **C:698** | 96 | SNP | | Salt_2_contig8611 | |
| **C:698** | 97 | SNP | | Salt_2_contig8611 | |
| **C:28635** | 84 | CHG | | Salt_2_contig9036 | |
| **C:21** | 135 | SNP | | Salt_2_contig9944 | |
| **C:1000** | 109 | SNP | | | |
| **C:1001** | 191 | SNP | | | |
| **C:1004** | 141 | SNP | | | |
| **C:10065** | 89 | SNP | | | |
| **C:10069** | 148 | SNP | | | |
| **C:10074** | 76 | CHG | | | |
| **C:10092** | 62 | SNP | | | |
| **C:10100** | 8 | SNP | | | |

| | | |
|---|---|---|
| **C:10112** | 147 | SNP |
| **C:10117** | 29 | SNP |
| **C:1012** | 114 | SNP |
| **C:10154** | 141 | SNP |
| **C:10192** | 129 | SNP |
| **C:10209** | 19 | SNP |
| **C:10209** | 90 | SNP |
| **C:10209** | 93 | SNP |
| **C:1277** | 151 | SNP |
| **C:128** | 174 | SNP |
| **C:10217** | 116 | SNP |
| **C:10219** | 192 | CG |
| **C:10234** | 42 | SNP |
| **C:12878** | 30 | SNP |
| **C:1291** | 58 | SNP |
| **C:1293** | 99 | SNP |
| **C:10243** | 83 | SNP |
| **C:10272** | 78 | SNP |
| **C:10278** | 149 | SNP |
| **C:1028** | 14 | SNP |
| **C:10303** | 157 | SNP |
| **C:10322** | 127 | SNP |
| **C:10414** | 58 | SNP |
| **C:1042** | 82 | SNP |
| **C:10421** | 78 | SNP |
| **C:10433** | 128 | SNP |
| **C:10433** | 158 | SNP |
| **C:10464** | 55 | SNP |
| **C:10491** | 96 | SNP |
| **C:105** | 39 | SNP |
| **C:10500** | 70 | SNP |
| **C:10567** | 42 | SNP |

| | | |
|---|---|---|
| **C:10567** | 49 | SNP |
| **C:10567** | 105 | SNP |
| **C:13576** | 26 | SNP |
| **C:136** | 73 | SNP |
| **C:13600** | 52 | SNP |
| **C:10604** | 41 | SNP |
| **C:10604** | 72 | SNP |
| **C:10669** | 10 | SNP |
| **C:10670** | 36 | SNP |
| **C:10684** | 128 | CHG |
| **C:107** | 10 | SNP |
| **C:107** | 89 | SNP |
| **C:107** | 145 | SNP |
| **C:107** | 166 | SNP |
| **C:107** | 184 | SNP |
| **C:107** | 55 | CHH |
| **C:10710** | 135 | SNP |
| **C:10713** | 153 | SNP |
| **C:10741** | 15 | SNP |
| **C:10781** | 51 | SNP |
| **C:10783** | 55 | SNP |
| **C:10783** | 87 | SNP |
| **C:10792** | 16 | SNP |
| **C:10814** | 77 | CHH |
| **C:10823** | 7 | SNP |
| **C:1084** | 144 | SNP |
| **C:1084** | 158 | SNP |
| **C:10852** | 51 | SNP |
| **C:10876** | 24 | SNP |
| **C:10880** | 55 | SNP |
| **C:10900** | 91 | SNP |
| **C:10911** | 116 | SNP |

| | | |
|---|---|---|
| **C:10919** | 102 | SNP |
| **C:1092** | 35 | SNP |
| **C:10920** | 75 | SNP |
| **C:1093** | 93 | CHH |
| **C:10938** | 150 | SNP |
| **C:10944** | 53 | SNP |
| **C:10944** | 119 | SNP |
| **C:10951** | 112 | SNP |
| **C:10951** | 79 | CHG |
| **C:10951** | 99 | CG |
| **C:10959** | 27 | SNP |
| **C:10962** | 45 | SNP |
| **C:10962** | 55 | SNP |
| **C:10962** | 62 | SNP |
| **C:1098** | 22 | SNP |
| **C:1098** | 26 | SNP |
| **C:10981** | 168 | CG |
| **C:10990** | 14 | SNP |
| **C:13994** | 79 | SNP |
| **C:14003** | 88 | CHH |
| **C:11022** | 32 | SNP |
| **C:11024** | 147 | SNP |
| **C:11039** | 60 | SNP |
| **C:1104** | 78 | SNP |
| **C:1105** | 16 | SNP |
| **C:11065** | 132 | CG |
| **C:11074** | 95 | SNP |
| **C:1108** | 10 | SNP |
| **C:11081** | 116 | SNP |
| **C:11083** | 95 | SNP |
| **C:11087** | 83 | SNP |
| **C:11090** | 80 | SNP |

| | | |
|---|---|---|
| **C:11090** | 94 | SNP |
| **C:11101** | 63 | SNP |
| **C:11101** | 121 | SNP |
| **C:11107** | 155 | SNP |
| **C:11107** | 67 | CHG |
| **C:11125** | 21 | SNP |
| **C:14057** | 37 | SNP |
| **C:11138** | 39 | CG |
| **C:11138** | 38 | CG |
| **C:1114** | 149 | SNP |
| **C:1114** | 155 | SNP |
| **C:11141** | 38 | CG |
| **C:11146** | 43 | SNP |
| **C:11146** | 55 | SNP |
| **C:11147** | 84 | SNP |
| **C:11147** | 86 | SNP |
| **C:1115** | 123 | SNP |
| **C:11154** | 13 | SNP |
| **C:14135** | 80 | SNP |
| **C:1414** | 36 | SNP |
| **C:14147** | 114 | SNP |
| **C:11166** | 87 | SNP |
| **C:11166** | 52 | SNP |
| **C:11182** | 25 | SNP |
| **C:11182** | 57 | SNP |
| **C:11182** | 92 | CHG |
| **C:11185** | 88 | CHH |
| **C:11187** | 17 | SNP |
| **C:11187** | 96 | SNP |
| **C:11189** | 98 | SNP |
| **C:11197** | 48 | SNP |
| **C:11205** | 135 | CHH |

| | | |
|---|---|---|
| **C:14237** | 9 | SNP |
| **C:11214** | 166 | SNP |
| **C:11221** | 71 | SNP |
| **C:11222** | 126 | CHH |
| **C:11226** | 70 | SNP |
| **C:11226** | 126 | SNP |
| **C:11226** | 129 | SNP |
| **C:11239** | 143 | SNP |
| **C:11242** | 29 | SNP |
| **C:11242** | 127 | SNP |
| **C:11255** | 133 | SNP |
| **C:11266** | 24 | SNP |
| **C:11267** | 112 | SNP |
| **C:11285** | 25 | SNP |
| **C:11286** | 168 | SNP |
| **C:11296** | 79 | SNP |
| **C:11296** | 121 | SNP |
| **C:11297** | 29 | SNP |
| **C:11297** | 107 | SNP |
| **C:11297** | 68 | SNP |
| **C:113** | 187 | SNP |
| **C:11301** | 46 | SNP |
| **C:11301** | 63 | SNP |
| **C:11303** | 42 | SNP |
| **C:11305** | 75 | SNP |
| **C:11305** | 95 | SNP |
| **C:11305** | 15 | SNP |
| **C:1131** | 130 | SNP |
| **C:11314** | 155 | CHH |
| **C:11318** | 80 | SNP |
| **C:11323** | 81 | SNP |
| **C:11326** | 130 | SNP |

| | | |
|---|---|---|
| **C:11328** | 118 | SNP |
| **C:11328** | 112 | SNP |
| **C:11328** | 120 | SNP |
| **C:11334** | 65 | SNP |
| **C:11335** | 117 | SNP |
| **C:11340** | 145 | SNP |
| **C:11340** | 85 | SNP |
| **C:11354** | 8 | SNP |
| **C:11354** | 109 | SNP |
| **C:14775** | 115 | SNP |
| **C:11378** | 147 | SNP |
| **C:11379** | 82 | SNP |
| **C:11381** | 35 | SNP |
| **C:11386** | 88 | CHH |
| **C:11389** | 150 | SNP |
| **C:114** | 21 | SNP |
| **C:11412** | 131 | SNP |
| **C:11415** | 99 | SNP |
| **C:11415** | 105 | SNP |
| **C:11418** | 24 | SNP |
| **C:1142** | 12 | CHG |
| **C:11422** | 140 | SNP |
| **C:11428** | 156 | CG |
| **C:11428** | 15 | CG |
| **C:11430** | 56 | SNP |
| **C:11443** | 126 | SNP |
| **C:11452** | 74 | SNP |
| **C:11461** | 91 | SNP |
| **C:11461** | 25 | SNP |
| **C:11464** | 53 | SNP |
| **C:11464** | 157 | SNP |
| **C:11467** | 74 | SNP |

| | | |
|---|---|---|
| **C:11472** | 163 | SNP |
| **C:11496** | 36 | SNP |
| **C:11496** | 118 | SNP |
| **C:11497** | 89 | CG |
| **C:11504** | 40 | SNP |
| **C:1151** | 95 | SNP |
| **C:11516** | 61 | SNP |
| **C:11516** | 136 | SNP |
| **C:11522** | 12 | SNP |
| **C:11522** | 160 | SNP |
| **C:11522** | 110 | SNP |
| **C:11527** | 22 | SNP |
| **C:15717** | 168 | SNP |
| **C:11534** | 127 | SNP |
| **C:11534** | 89 | SNP |
| **C:11538** | 142 | SNP |
| **C:11554** | 31 | SNP |
| **C:11562** | 79 | SNP |
| **C:11569** | 60 | SNP |
| **C:11590** | 44 | SNP |
| **C:11591** | 53 | SNP |
| **C:11591** | 121 | SNP |
| **C:11614** | 94 | SNP |
| **C:11623** | 18 | CG |
| **C:11630** | 75 | SNP |
| **C:11630** | 90 | SNP |
| **C:11643** | 143 | CG |
| **C:1165** | 15 | CHH |
| **C:11653** | 60 | SNP |
| **C:11655** | 125 | SNP |
| **C:1166** | 30 | SNP |
| **C:11660** | 6 | SNP |

| | | |
|---|---|---|
| **C:11662** | 17 | SNP |
| **C:11670** | 45 | SNP |
| **C:11680** | 86 | SNP |
| **C:11683** | 147 | SNP |
| **C:11684** | 104 | SNP |
| **C:11699** | 14 | SNP |
| **C:11705** | 93 | SNP |
| **C:1171** | 37 | SNP |
| **C:1171** | 38 | SNP |
| **C:11711** | 137 | SNP |
| **C:11714** | 109 | SNP |
| **C:11726** | 33 | SNP |
| **C:11733** | 129 | SNP |
| **C:11735** | 41 | SNP |
| **C:11738** | 142 | SNP |
| **C:11741** | 148 | SNP |
| **C:11741** | 51 | SNP |
| **C:11741** | 102 | SNP |
| **C:11742** | 99 | SNP |
| **C:11754** | 139 | SNP |
| **C:11754** | 151 | SNP |
| **C:11754** | 102 | SNP |
| **C:11756** | 146 | SNP |
| **C:11759** | 97 | SNP |
| **C:11764** | 93 | SNP |
| **C:11768** | 37 | SNP |
| **C:11778** | 24 | SNP |
| **C:11778** | 34 | SNP |
| **C:11778** | 55 | SNP |
| **C:11778** | 160 | SNP |
| **C:11783** | 25 | SNP |
| **C:11783** | 48 | SNP |

| | | |
|---|---|---|
| **C:11787** | 160 | SNP |
| **C:11796** | 150 | SNP |
| **C:11802** | 98 | SNP |
| **C:11806** | 88 | SNP |
| **C:11815** | 157 | SNP |
| **C:11815** | 27 | SNP |
| **C:1183** | 115 | SNP |
| **C:11838** | 18 | SNP |
| **C:11838** | 95 | SNP |
| **C:1829** | 153 | SNP |
| **C:11847** | 147 | SNP |
| **C:11847** | 162 | SNP |
| **C:11847** | 173 | SNP |
| **C:11853** | 140 | SNP |
| **C:11854** | 99 | SNP |
| **C:11854** | 117 | SNP |
| **C:1186** | 128 | SNP |
| **C:1187** | 84 | SNP |
| **C:11877** | 76 | SNP |
| **C:11879** | 117 | SNP |
| **C:1188** | 168 | SNP |
| **C:11889** | 85 | SNP |
| **C:11891** | 11 | SNP |
| **C:11891** | 152 | SNP |
| **C:11901** | 28 | SNP |
| **C:11919** | 40 | SNP |
| **C:11927** | 52 | SNP |
| **C:11930** | 88 | SNP |
| **C:11930** | 94 | CHG |
| **C:11931** | 60 | SNP |
| **C:11943** | 149 | SNP |
| **C:11943** | 148 | SNP |

| | | |
|---|---|---|
| **C:11944** | 17 | SNP |
| **C:11949** | 25 | SNP |
| **C:1195** | 72 | SNP |
| **C:11950** | 141 | SNP |
| **C:11950** | 18 | SNP |
| **C:11950** | 122 | SNP |
| **C:11958** | 73 | SNP |
| **C:11967** | 88 | SNP |
| **C:12023** | 47 | SNP |
| **C:12023** | 83 | SNP |
| **C:12024** | 34 | SNP |
| **C:12024** | 67 | SNP |
| **C:12029** | 60 | SNP |
| **C:1203** | 117 | SNP |
| **C:1203** | 129 | CHG |
| **C:12034** | 81 | SNP |
| **C:1205** | 16 | CG |
| **C:1207** | 172 | SNP |
| **C:12102** | 47 | SNP |
| **C:1211** | 35 | SNP |
| **C:1213** | 50 | SNP |
| **C:1217** | 49 | SNP |
| **C:12192** | 69 | SNP |
| **C:122** | 173 | SNP |
| **C:12216** | 16 | SNP |
| **C:1224** | 187 | SNP |
| **C:2054** | 139 | SNP |
| **C:12298** | 86 | CG |
| **C:12302** | 38 | SNP |
| **C:12359** | 87 | CG |
| **C:12394** | 71 | SNP |
| **C:1241** | 47 | SNP |

| | | |
|---|---|---|
| **C:12424** | 49 | SNP |
| **C:1243** | 43 | SNP |
| **C:12433** | 94 | SNP |
| **C:12456** | 81 | SNP |
| **C:12468** | 110 | SNP |
| **C:1249** | 58 | SNP |
| **C:1249** | 134 | SNP |
| **C:12498** | 23 | SNP |
| **C:12498** | 47 | SNP |
| **C:12498** | 110 | SNP |
| **C:2113** | 36 | SNP |
| **C:12516** | 18 | CG |
| **C:2117** | 192 | SNP |
| **C:12533** | 52 | SNP |
| **C:1254** | 19 | SNP |
| **C:1254** | 86 | SNP |
| **C:12548** | 27 | SNP |
| **C:12548** | 75 | SNP |
| **C:12569** | 48 | SNP |
| **C:1260** | 25 | SNP |
| **C:1260** | 62 | CHH |
| **C:1261** | 97 | SNP |
| **C:12613** | 12 | SNP |
| **C:12613** | 88 | SNP |
| **C:1262** | 196 | SNP |
| **C:12629** | 78 | SNP |
| **C:12649** | 65 | SNP |
| **C:12650** | 81 | SNP |
| **C:12670** | 29 | SNP |
| **C:12677** | 43 | CHG |
| **C:2256** | 136 | SNP |
| **C:226** | 33 | SNP |

| | | |
|---|---|---|
| **C:226** | 68 | SNP |
| **C:12837** | 37 | SNP |
| **C:12837** | 59 | SNP |
| **C:1284** | 13 | SNP |
| **C:1284** | 170 | SNP |
| **C:2292** | 57 | SNP |
| **C:2292** | 93 | CG |
| **C:2298** | 28 | SNP |
| **C:34244** | 151 | SNP |
| **C:13146** | 37 | SNP |
| **C:1316** | 169 | SNP |
| **C:13227** | 50 | SNP |
| **C:1325** | 76 | SNP |
| **C:1327** | 62 | SNP |
| **C:1327** | 122 | SNP |
| **C:13294** | 51 | SNP |
| **C:133** | 33 | SNP |
| **C:134** | 192 | SNP |
| **C:13502** | 133 | SNP |
| **C:13504** | 120 | CG |
| **C:13519** | 57 | CHG |
| **C:13520** | 125 | CHH |
| **C:13537** | 14 | SNP |
| **C:13540** | 99 | SNP |
| **C:13543** | 112 | CHH |
| **C:13546** | 130 | SNP |
| **C:13550** | 15 | SNP |
| **C:2439** | 145 | SNP |
| **C:13601** | 28 | SNP |
| **C:13629** | 20 | SNP |
| **C:13648** | 74 | SNP |
| **C:13672** | 15 | SNP |

| | | |
|---|---|---|
| **C:13672** | 21 | SNP |
| **C:13672** | 123 | SNP |
| **C:13673** | 58 | SNP |
| **C:13681** | 16 | SNP |
| **C:137** | 178 | SNP |
| **C:13706** | 113 | SNP |
| **C:13713** | 63 | SNP |
| **C:13717** | 58 | SNP |
| **C:13726** | 17 | SNP |
| **C:2520** | 153 | CG |
| **C:253** | 30 | SNP |
| **C:253** | 133 | SNP |
| **C:2561** | 13 | SNP |
| **C:13772** | 131 | SNP |
| **C:13773** | 66 | SNP |
| **C:13784** | 131 | SNP |
| **C:13810** | 82 | SNP |
| **C:13820** | 50 | CG |
| **C:13820** | 129 | CHG |
| **C:13820** | 38 | CG |
| **C:13826** | 128 | SNP |
| **C:13836** | 14 | SNP |
| **C:13836** | 49 | SNP |
| **C:13842** | 11 | SNP |
| **C:13857** | 70 | SNP |
| **C:13859** | 107 | SNP |
| **C:13865** | 13 | CG |
| **C:13865** | 11 | CG |
| **C:13868** | 20 | CHH |
| **C:2667** | 137 | SNP |
| **C:13899** | 81 | SNP |
| **C:139** | 168 | SNP |

| | | |
|---|---|---|
| **C:13901** | 92 | SNP |
| **C:13901** | 120 | SNP |
| **C:13910** | 24 | SNP |
| **C:13910** | 45 | SNP |
| **C:13920** | 92 | SNP |
| **C:13920** | 61 | CG |
| **C:13920** | 60 | CG |
| **C:13931** | 78 | SNP |
| **C:13947** | 23 | SNP |
| **C:13976** | 101 | SNP |
| **C:13977** | 97 | CHG |
| **C:13991** | 66 | SNP |
| **C:2681** | 90 | SNP |
| **C:2681** | 62 | SNP |
| **C:14004** | 61 | SNP |
| **C:14006** | 141 | CHH |
| **C:2687** | 13 | SNP |
| **C:2690** | 28 | SNP |
| **C:2690** | 89 | SNP |
| **C:26956** | 67 | SNP |
| **C:14025** | 136 | CHH |
| **C:14031** | 140 | CG |
| **C:14034** | 58 | SNP |
| **C:14035** | 84 | SNP |
| **C:27092** | 47 | SNP |
| **C:14045** | 39 | SNP |
| **C:14048** | 119 | SNP |
| **C:14049** | 138 | SNP |
| **C:14049** | 90 | SNP |
| **C:14049** | 106 | SNP |
| **C:14051** | 20 | SNP |
| **C:27264** | 118 | SNP |

| | | |
|---|---|---|
| **C:14069** | 23 | SNP |
| **C:1407** | 14 | SNP |
| **C:1407** | 58 | SNP |
| **C:14076** | 96 | CHH |
| **C:14083** | 73 | SNP |
| **C:14083** | 127 | SNP |
| **C:14085** | 77 | SNP |
| **C:14085** | 103 | SNP |
| **C:1411** | 178 | SNP |
| **C:14111** | 91 | SNP |
| **C:14128** | 90 | SNP |
| **C:27442** | 30 | SNP |
| **C:27459** | 116 | SNP |
| **C:27481** | 98 | SNP |
| **C:14152** | 75 | SNP |
| **C:14156** | 138 | SNP |
| **C:14161** | 131 | CHH |
| **C:14172** | 139 | SNP |
| **C:14172** | 129 | SNP |
| **C:14176** | 124 | SNP |
| **C:14176** | 134 | SNP |
| **C:14190** | 28 | SNP |
| **C:14230** | 132 | SNP |
| **C:14230** | 81 | CHH |
| **C:14237** | 83 | SNP |
| **C:27738** | 31 | SNP |
| **C:14237** | 81 | SNP |
| **C:14264** | 62 | SNP |
| **C:14269** | 52 | SNP |
| **C:14269** | 97 | SNP |
| **C:1427** | 166 | SNP |
| **C:14311** | 21 | SNP |

| | | |
|---|---|---|
| **C:14313** | 19 | SNP |
| **C:14314** | 109 | SNP |
| **C:14314** | 112 | SNP |
| **C:14331** | 113 | SNP |
| **C:1434** | 33 | SNP |
| **C:1434** | 179 | SNP |
| **C:1434** | 178 | CG |
| **C:14348** | 26 | SNP |
| **C:28065** | 123 | CG |
| **C:14408** | 38 | SNP |
| **C:14414** | 93 | SNP |
| **C:14414** | 118 | SNP |
| **C:14551** | 45 | SNP |
| **C:14567** | 77 | SNP |
| **C:14567** | 51 | SNP |
| **C:14568** | 37 | SNP |
| **C:14568** | 44 | SNP |
| **C:1459** | 182 | SNP |
| **C:1459** | 108 | SNP |
| **C:14639** | 25 | SNP |
| **C:14639** | 78 | SNP |
| **C:14639** | 98 | SNP |
| **C:14639** | 103 | SNP |
| **C:14658** | 117 | SNP |
| **C:14662** | 47 | SNP |
| **C:14710** | 76 | SNP |
| **C:14719** | 56 | SNP |
| **C:14719** | 34 | CG |
| **C:14719** | 11 | SNP |
| **C:14719** | 98 | SNP |
| **C:14722** | 9 | SNP |
| **C:14722** | 74 | SNP |

| | | |
|---|---|---|
| **C:1473** | 123 | SNP |
| **C:1473** | 129 | SNP |
| **C:14773** | 86 | SNP |
| **C:28724** | 14 | SNP |
| **C:14775** | 96 | SNP |
| **C:14780** | 64 | SNP |
| **C:14781** | 100 | CHH |
| **C:14804** | 93 | SNP |
| **C:14807** | 115 | SNP |
| **C:14811** | 40 | SNP |
| **C:14836** | 77 | SNP |
| **C:14862** | 57 | CG |
| **C:14879** | 18 | SNP |
| **C:14903** | 105 | SNP |
| **C:14908** | 29 | SNP |
| **C:1491** | 90 | SNP |
| **C:14959** | 110 | SNP |
| **C:1498** | 114 | CHG |
| **C:14990** | 65 | SNP |
| **C:14990** | 110 | SNP |
| **C:150** | 179 | SNP |
| **C:15001** | 28 | SNP |
| **C:15005** | 54 | SNP |
| **C:15005** | 96 | SNP |
| **C:15005** | 51 | SNP |
| **C:15005** | 111 | SNP |
| **C:1503** | 40 | SNP |
| **C:1504** | 49 | SNP |
| **C:1515** | 53 | CG |
| **C:15156** | 41 | SNP |
| **C:1529** | 151 | SNP |
| **C:15427** | 26 | SNP |

| | | |
|---|---|---|
| **C:1543** | 128 | SNP |
| **C:1543** | 165 | SNP |
| **C:1543** | 186 | SNP |
| **C:295** | 154 | SNP |
| **C:1560** | 24 | SNP |
| **C:1565** | 177 | SNP |
| **C:1567** | 40 | SNP |
| **C:29557** | 90 | SNP |
| **C:1577** | 142 | SNP |
| **C:158** | 67 | SNP |
| **C:1592** | 26 | SNP |
| **C:1598** | 6 | SNP |
| **C:2964** | 28 | SNP |
| **C:1605** | 162 | SNP |
| **C:16112** | 144 | SNP |
| **C:16112** | 143 | SNP |
| **C:1613** | 158 | SNP |
| **C:1613** | 177 | SNP |
| **C:1614** | 165 | SNP |
| **C:16167** | 160 | SNP |
| **C:3002** | 50 | SNP |
| **C:1630** | 54 | SNP |
| **C:1636** | 196 | SNP |
| **C:16404** | 189 | SNP |
| **C:16404** | 193 | SNP |
| **C:16404** | 145 | SNP |
| **C:1643** | 8 | SNP |
| **C:1652** | 47 | SNP |
| **C:16523** | 34 | SNP |
| **C:16533** | 64 | SNP |
| **C:16533** | 73 | SNP |
| **C:1659** | 194 | SNP |

| | | |
|---|---|---|
| **C:1678** | 136 | SNP |
| **C:168** | 25 | SNP |
| **C:1683** | 32 | SNP |
| **C:1683** | 184 | SNP |
| **C:1689** | 73 | SNP |
| **C:1695** | 186 | SNP |
| **C:17** | 109 | SNP |
| **C:170** | 29 | SNP |
| **C:17018** | 27 | SNP |
| **C:17108** | 177 | SNP |
| **C:1713** | 147 | SNP |
| **C:17139** | 13 | SNP |
| **C:172** | 32 | SNP |
| **C:17244** | 46 | SNP |
| **C:174** | 52 | SNP |
| **C:1741** | 91 | SNP |
| **C:1741** | 92 | SNP |
| **C:1746** | 53 | SNP |
| **C:1749** | 24 | SNP |
| **C:1756** | 185 | SNP |
| **C:1766** | 22 | SNP |
| **C:1766** | 21 | CG |
| **C:1768** | 22 | SNP |
| **C:1775** | 123 | SNP |
| **C:1776** | 33 | CHG |
| **C:178** | 128 | SNP |
| **C:179** | 74 | SNP |
| **C:1793** | 42 | SNP |
| **C:1797** | 50 | SNP |
| **C:1797** | 181 | SNP |
| **C:18** | 24 | SNP |
| **C:180** | 158 | SNP |

| | | |
|---|---|---|
| **C:1800** | 144 | SNP |
| **C:31128** | 90 | CG |
| **C:3123** | 117 | SNP |
| **C:1823** | 197 | SNP |
| **C:1824** | 179 | SNP |
| **C:1827** | 141 | CHH |
| **C:31268** | 37 | SNP |
| **C:1835** | 81 | SNP |
| **C:1835** | 174 | SNP |
| **C:18443** | 9 | SNP |
| **C:18443** | 71 | SNP |
| **C:1853** | 119 | CHG |
| **C:1874** | 72 | SNP |
| **C:1875** | 153 | SNP |
| **C:19** | 109 | SNP |
| **C:316** | 26 | SNP |
| **C:3166** | 193 | SNP |
| **C:31664** | 35 | SNP |
| **C:31686** | 57 | SNP |
| **C:191** | 8 | SNP |
| **C:191** | 67 | SNP |
| **C:1928** | 67 | SNP |
| **C:1956** | 68 | SNP |
| **C:1957** | 139 | SNP |
| **C:1957** | 153 | SNP |
| **C:1957** | 196 | SNP |
| **C:1960** | 93 | SNP |
| **C:1960** | 94 | SNP |
| **C:19620** | 45 | SNP |
| **C:19632** | 65 | SNP |
| **C:19632** | 84 | SNP |
| **C:1974** | 117 | SNP |

| | | |
|---|---|---|
| **C:1974** | 148 | SNP |
| **C:1974** | 171 | SNP |
| **C:1974** | 188 | SNP |
| **C:32058** | 29 | SNP |
| **C:32058** | 76 | SNP |
| **C:198** | 57 | SNP |
| **C:1983** | 58 | CHH |
| **C:1985** | 29 | SNP |
| **C:1994** | 91 | SNP |
| **C:1994** | 149 | CHG |
| **C:200** | 45 | SNP |
| **C:2003** | 44 | SNP |
| **C:2008** | 159 | SNP |
| **C:202** | 116 | SNP |
| **C:203** | 65 | SNP |
| **C:20353** | 60 | SNP |
| **C:2036** | 56 | SNP |
| **C:20376** | 72 | SNP |
| **C:204** | 161 | SNP |
| **C:2045** | 20 | SNP |
| **C:2045** | 35 | SNP |
| **C:205** | 69 | SNP |
| **C:32925** | 30 | SNP |
| **C:32925** | 61 | SNP |
| **C:32925** | 81 | SNP |
| **C:2058** | 132 | SNP |
| **C:20589** | 14 | SNP |
| **C:2060** | 21 | SNP |
| **C:2067** | 14 | SNP |
| **C:2070** | 191 | SNP |
| **C:2076** | 185 | SNP |
| **C:2093** | 44 | SNP |

| | | |
|---|---|---|
| **C:2094** | 51 | SNP |
| **C:2103** | 110 | SNP |
| **C:2103** | 64 | SNP |
| **C:2105** | 45 | SNP |
| **C:2109** | 92 | SNP |
| **C:2109** | 17 | SNP |
| **C:2113** | 178 | SNP |
| **C:33281** | 20 | SNP |
| **C:2117** | 171 | SNP |
| **C:3329** | 28 | SNP |
| **C:2122** | 43 | CG |
| **C:21238** | 146 | SNP |
| **C:2132** | 40 | SNP |
| **C:2141** | 185 | SNP |
| **C:2163** | 86 | SNP |
| **C:2173** | 9 | SNP |
| **C:2175** | 75 | SNP |
| **C:2175** | 103 | SNP |
| **C:218** | 85 | SNP |
| **C:2195** | 17 | SNP |
| **C:221** | 43 | SNP |
| **C:2212** | 132 | CHH |
| **C:2216** | 80 | SNP |
| **C:2245** | 87 | SNP |
| **C:2245** | 154 | SNP |
| **C:225** | 18 | SNP |
| **C:225** | 145 | SNP |
| **C:225** | 153 | CG |
| **C:340** | 108 | SNP |
| **C:34011** | 140 | SNP |
| **C:34011** | 148 | SNP |
| **C:2263** | 21 | CG |

| | | |
|---|---|---|
| **C:228** | 67 | SNP |
| **C:2286** | 108 | SNP |
| **C:2291** | 142 | SNP |
| **C:34129** | 122 | CG |
| **C:34169** | 38 | SNP |
| **C:34204** | 69 | CHH |
| **C:5620** | 13 | SNP |
| **C:2327** | 51 | SNP |
| **C:2330** | 127 | SNP |
| **C:2337** | 91 | SNP |
| **C:235** | 7 | SNP |
| **C:235** | 168 | SNP |
| **C:235** | 178 | SNP |
| **C:235** | 188 | SNP |
| **C:2352** | 195 | SNP |
| **C:236** | 180 | SNP |
| **C:2383** | 27 | SNP |
| **C:2383** | 104 | SNP |
| **C:2383** | 51 | CG |
| **C:2384** | 59 | SNP |
| **C:2384** | 185 | SNP |
| **C:2385** | 106 | SNP |
| **C:2399** | 112 | SNP |
| **C:2407** | 153 | SNP |
| **C:2408** | 67 | SNP |
| **C:34629** | 48 | SNP |
| **C:34632** | 43 | CHH |
| **C:3468** | 128 | SNP |
| **C:245** | 108 | SNP |
| **C:245** | 31 | CG |
| **C:247** | 188 | SNP |
| **C:2477** | 29 | SNP |

| | | |
|---|---|---|
| **C:24815** | 8 | SNP |
| **C:2482** | 16 | SNP |
| **C:25** | 15 | SNP |
| **C:25** | 116 | SNP |
| **C:2512** | 32 | SNP |
| **C:2512** | 74 | SNP |
| **C:252** | 177 | SNP |
| **C:2520** | 109 | SNP |
| **C:349** | 59 | CG |
| **C:3493** | 149 | CG |
| **C:34949** | 80 | SNP |
| **C:610** | 39 | SNP |
| **C:2562** | 98 | SNP |
| **C:257** | 83 | SNP |
| **C:257** | 110 | SNP |
| **C:257** | 152 | SNP |
| **C:2585** | 136 | SNP |
| **C:2593** | 46 | SNP |
| **C:2619** | 115 | SNP |
| **C:26405** | 118 | SNP |
| **C:35116** | 32 | SNP |
| **C:26514** | 64 | SNP |
| **C:2655** | 9 | SNP |
| **C:26582** | 30 | SNP |
| **C:26652** | 60 | SNP |
| **C:26652** | 154 | SNP |
| **C:2667** | 71 | SNP |
| **C:35357** | 55 | SNP |
| **C:2667** | 185 | SNP |
| **C:2668** | 63 | SNP |
| **C:26681** | 122 | SNP |
| **C:267** | 11 | SNP |

| | | |
|---|---|---|
| **C:267** | 19 | SNP |
| **C:2670** | 56 | SNP |
| **C:2672** | 49 | SNP |
| **C:26721** | 10 | CG |
| **C:26753** | 101 | SNP |
| **C:26760** | 105 | SNP |
| **C:26763** | 145 | SNP |
| **C:26768** | 8 | SNP |
| **C:26799** | 74 | SNP |
| **C:26805** | 111 | SNP |
| **C:35725** | 9 | SNP |
| **C:3579** | 87 | SNP |
| **C:2682** | 131 | SNP |
| **C:26833** | 114 | SNP |
| **C:3582** | 37 | SNP |
| **C:3582** | 109 | SNP |
| **C:35827** | 35 | CG |
| **C:6565** | 94 | CG |
| **C:35918** | 41 | CG |
| **C:26993** | 134 | CG |
| **C:35930** | 83 | SNP |
| **C:36090** | 126 | SNP |
| **C:27092** | 20 | SNP |
| **C:3616** | 83 | SNP |
| **C:27092** | 49 | SNP |
| **C:27116** | 145 | SNP |
| **C:27147** | 21 | CG |
| **C:27165** | 94 | CG |
| **C:27167** | 86 | CHG |
| **C:27246** | 27 | SNP |
| **C:363** | 10 | SNP |
| **C:27294** | 122 | SNP |

| | | |
|---|---|---|
| **C:27330** | 34 | SNP |
| **C:2734** | 137 | SNP |
| **C:27356** | 23 | SNP |
| **C:27358** | 133 | SNP |
| **C:3674** | 147 | SNP |
| **C:274** | 38 | SNP |
| **C:27420** | 16 | SNP |
| **C:27428** | 10 | SNP |
| **C:27428** | 19 | SNP |
| **C:27430** | 120 | SNP |
| **C:6708** | 95 | SNP |
| **C:6714** | 117 | CHG |
| **C:3710** | 44 | SNP |
| **C:27514** | 74 | SNP |
| **C:27514** | 16 | CHG |
| **C:276** | 64 | SNP |
| **C:2761** | 71 | SNP |
| **C:27661** | 25 | SNP |
| **C:27661** | 26 | SNP |
| **C:27670** | 84 | SNP |
| **C:27714** | 124 | SNP |
| **C:27726** | 97 | SNP |
| **C:27726** | 108 | SNP |
| **C:27758** | 144 | SNP |
| **C:27794** | 123 | SNP |
| **C:278** | 122 | SNP |
| **C:3769** | 123 | SNP |
| **C:2793** | 93 | SNP |
| **C:27962** | 13 | SNP |
| **C:27962** | 137 | SNP |
| **C:27984** | 91 | SNP |
| **C:27989** | 74 | SNP |

| | | |
|---|---|---|
| **C:28** | 106 | SNP |
| **C:28061** | 10 | SNP |
| **C:28061** | 37 | SNP |
| **C:28061** | 49 | SNP |
| **C:28061** | 58 | SNP |
| **C:28061** | 63 | SNP |
| **C:3859** | 62 | SNP |
| **C:2808** | 153 | SNP |
| **C:2810** | 64 | SNP |
| **C:2810** | 55 | SNP |
| **C:28179** | 99 | SNP |
| **C:2827** | 25 | SNP |
| **C:2827** | 49 | SNP |
| **C:2827** | 109 | SNP |
| **C:28307** | 91 | SNP |
| **C:28316** | 112 | SNP |
| **C:28369** | 117 | SNP |
| **C:28370** | 10 | SNP |
| **C:28380** | 118 | SNP |
| **C:28467** | 86 | SNP |
| **C:285** | 88 | CG |
| **C:285** | 36 | CG |
| **C:285** | 89 | CG |
| **C:2852** | 145 | SNP |
| **C:2852** | 114 | SNP |
| **C:28537** | 86 | SNP |
| **C:28571** | 60 | SNP |
| **C:28596** | 88 | SNP |
| **C:28622** | 43 | CG |
| **C:28651** | 77 | SNP |
| **C:287** | 113 | SNP |
| **C:4049** | 15 | SNP |

| | | |
|---|---|---|
| **C:28734** | 143 | CG |
| **C:2874** | 76 | SNP |
| **C:28770** | 62 | SNP |
| **C:288** | 9 | SNP |
| **C:28941** | 25 | CHG |
| **C:2897** | 76 | SNP |
| **C:29** | 21 | SNP |
| **C:29** | 80 | SNP |
| **C:29083** | 68 | CHG |
| **C:29089** | 10 | SNP |
| **C:29124** | 34 | CHH |
| **C:2913** | 70 | SNP |
| **C:29135** | 47 | SNP |
| **C:29212** | 75 | SNP |
| **C:29214** | 169 | CHG |
| **C:29214** | 174 | CHG |
| **C:29214** | 64 | CG |
| **C:29216** | 18 | SNP |
| **C:29216** | 125 | SNP |
| **C:29217** | 128 | SNP |
| **C:29241** | 77 | SNP |
| **C:2925** | 190 | SNP |
| **C:2927** | 60 | SNP |
| **C:2927** | 78 | SNP |
| **C:2928** | 16 | SNP |
| **C:29289** | 120 | SNP |
| **C:29394** | 99 | SNP |
| **C:4271** | 128 | SNP |
| **C:295** | 173 | SNP |
| **C:29519** | 172 | SNP |
| **C:29546** | 179 | CHH |
| **C:4290** | 31 | SNP |

| | | |
|---|---|---|
| **C:29564** | 76 | SNP |
| **C:29564** | 74 | SNP |
| **C:2958** | 178 | SNP |
| **C:2959** | 75 | SNP |
| **C:2964** | 47 | SNP |
| **C:29878** | 130 | SNP |
| **C:29878** | 166 | SNP |
| **C:299** | 63 | SNP |
| **C:299** | 89 | SNP |
| **C:2996** | 49 | SNP |
| **C:3** | 11 | SNP |
| **C:4386** | 36 | SNP |
| **C:30040** | 153 | SNP |
| **C:4415** | 97 | SNP |
| **C:30099** | 49 | SNP |
| **C:30102** | 53 | SNP |
| **C:30145** | 165 | SNP |
| **C:30145** | 77 | CHG |
| **C:30148** | 7 | SNP |
| **C:30174** | 93 | SNP |
| **C:4492** | 30 | SNP |
| **C:30298** | 94 | SNP |
| **C:45** | 158 | SNP |
| **C:45** | 27 | CHH |
| **C:30316** | 159 | CHG |
| **C:304** | 14 | SNP |
| **C:3042** | 42 | SNP |
| **C:4548** | 192 | SNP |
| **C:455** | 57 | SNP |
| **C:4561** | 76 | SNP |
| **C:46** | 154 | SNP |
| **C:30505** | 129 | SNP |

| | | |
|---|---|---|
| **C:30530** | 137 | CHH |
| **C:30563** | 158 | SNP |
| **C:30578** | 78 | SNP |
| **C:3058** | 60 | SNP |
| **C:3060** | 90 | SNP |
| **C:3060** | 131 | SNP |
| **C:30606** | 120 | SNP |
| **C:30666** | 77 | SNP |
| **C:30666** | 83 | SNP |
| **C:30669** | 139 | SNP |
| **C:307** | 41 | SNP |
| **C:30770** | 120 | SNP |
| **C:30798** | 15 | SNP |
| **C:30798** | 58 | CHG |
| **C:308** | 54 | SNP |
| **C:30824** | 68 | SNP |
| **C:30862** | 92 | CHH |
| **C:30898** | 76 | SNP |
| **C:30996** | 156 | CG |
| **C:3105** | 26 | SNP |
| **C:31067** | 82 | SNP |
| **C:31095** | 59 | SNP |
| **C:31103** | 131 | SNP |
| **C:31108** | 167 | SNP |
| **C:832** | 63 | SNP |
| **C:8340** | 151 | SNP |
| **C:3123** | 123 | CHG |
| **C:31231** | 15 | SNP |
| **C:31235** | 169 | CHH |
| **C:480** | 36 | SNP |
| **C:8387** | 138 | SNP |
| **C:3133** | 141 | SNP |

| | | |
|---|---|---|
| **C:3133** | 146 | SNP |
| **C:314** | 9 | SNP |
| **C:31483** | 112 | SNP |
| **C:315** | 62 | SNP |
| **C:315** | 66 | SNP |
| **C:315** | 70 | SNP |
| **C:315** | 89 | SNP |
| **C:4881** | 31 | SNP |
| **C:4893** | 120 | CG |
| **C:8592** | 12 | SNP |
| **C:4911** | 121 | SNP |
| **C:31722** | 8 | SNP |
| **C:31722** | 44 | SNP |
| **C:3182** | 52 | SNP |
| **C:3182** | 87 | SNP |
| **C:3182** | 184 | SNP |
| **C:31855** | 89 | SNP |
| **C:31883** | 8 | SNP |
| **C:31888** | 19 | SNP |
| **C:3189** | 163 | SNP |
| **C:31890** | 37 | SNP |
| **C:31908** | 86 | SNP |
| **C:31911** | 93 | CHH |
| **C:31922** | 20 | SNP |
| **C:31924** | 46 | SNP |
| **C:3194** | 179 | SNP |
| **C:32011** | 192 | CG |
| **C:8737** | 140 | SNP |
| **C:32087** | 22 | SNP |
| **C:32127** | 6 | SNP |
| **C:32128** | 9 | SNP |
| **C:32203** | 92 | SNP |

| | | |
|---|---|---|
| **C:32203** | 21 | SNP |
| **C:3225** | 53 | SNP |
| **C:323** | 120 | SNP |
| **C:323** | 146 | SNP |
| **C:32313** | 187 | CHG |
| **C:32338** | 80 | SNP |
| **C:3239** | 21 | SNP |
| **C:3241** | 79 | SNP |
| **C:32492** | 20 | SNP |
| **C:32582** | 55 | SNP |
| **C:326** | 21 | SNP |
| **C:32624** | 81 | SNP |
| **C:3266** | 108 | SNP |
| **C:32910** | 92 | SNP |
| **C:52** | 191 | SNP |
| **C:521** | 63 | SNP |
| **C:521** | 196 | SNP |
| **C:3297** | 27 | SNP |
| **C:3298** | 41 | SNP |
| **C:3298** | 100 | SNP |
| **C:3299** | 189 | SNP |
| **C:3299** | 69 | CG |
| **C:33** | 71 | CHG |
| **C:331** | 197 | SNP |
| **C:3311** | 8 | SNP |
| **C:3312** | 17 | SNP |
| **C:3312** | 27 | SNP |
| **C:3313** | 28 | CHH |
| **C:3317** | 165 | SNP |
| **C:33208** | 105 | SNP |
| **C:33269** | 49 | . |
| **C:33269** | 50 | SNP |

| | | |
|---|---|---|
| **C:535** | 38 | SNP |
| **C:33281** | 50 | SNP |
| **C:5355** | 140 | SNP |
| **C:3338** | 169 | CHH |
| **C:334** | 78 | SNP |
| **C:3351** | 91 | SNP |
| **C:3351** | 108 | SNP |
| **C:3351** | 109 | SNP |
| **C:3351** | 110 | SNP |
| **C:33599** | 102 | SNP |
| **C:33733** | 10 | SNP |
| **C:5390** | 130 | SNP |
| **C:33828** | 36 | SNP |
| **C:33861** | 48 | SNP |
| **C:33914** | 125 | CHG |
| **C:33995** | 44 | SNP |
| **C:33995** | 86 | SNP |
| **C:33995** | 105 | SNP |
| **C:34** | 70 | SNP |
| **C:34** | 89 | SNP |
| **C:340** | 106 | SNP |
| **C:5531** | 49 | SNP |
| **C:5531** | 121 | SNP |
| **C:554** | 88 | SNP |
| **C:34011** | 176 | SNP |
| **C:3410** | 8 | SNP |
| **C:3410** | 106 | SNP |
| **C:5577** | 113 | SNP |
| **C:559** | 130 | SNP |
| **C:5597** | 88 | SNP |
| **C:9738** | 123 | SNP |
| **C:34244** | 181 | SNP |

| | | |
|---|---|---|
| **C:34268** | 54 | SNP |
| **C:34292** | 48 | SNP |
| **C:34318** | 47 | SNP |
| **C:34407** | 33 | SNP |
| **C:345** | 96 | SNP |
| **C:345** | 185 | CHG |
| **C:34522** | 56 | SNP |
| **C:34539** | 7 | SNP |
| **C:34539** | 111 | SNP |
| **C:34569** | 123 | SNP |
| **C:3457** | 132 | SNP |
| **C:3457** | 150 | SNP |
| **C:34574** | 84 | SNP |
| **C:34574** | 71 | SNP |
| **C:34578** | 43 | SNP |
| **C:34586** | 91 | SNP |
| **C:34590** | 68 | SNP |
| **C:5959** | 18 | CG |
| **C:5961** | 136 | SNP |
| **C:5980** | 24 | SNP |
| **C:34711** | 47 | CHG |
| **C:34718** | 76 | SNP |
| **C:34732** | 56 | SNP |
| **C:34762** | 120 | SNP |
| **C:34818** | 107 | SNP |
| **C:34824** | 111 | SNP |
| **C:34824** | 139 | SNP |
| **C:34826** | 77 | SNP |
| **C:34851** | 19 | SNP |
| **C:34851** | 133 | SNP |
| **C:34894** | 68 | SNP |
| **C:34898** | 39 | CG |

| | | |
|---|---|---|
| **C:349** | 73 | SNP |
| **C:6064** | 87 | SNP |
| **C:6074** | 50 | SNP |
| **C:6074** | 123 | SNP |
| **C:4008** | 191 | SNP |
| **C:35004** | 71 | SNP |
| **C:35004** | 112 | SNP |
| **C:35018** | 79 | CG |
| **C:35041** | 58 | SNP |
| **C:35092** | 26 | SNP |
| **C:35092** | 66 | SNP |
| **C:3511** | 74 | SNP |
| **C:35116** | 67 | SNP |
| **C:623** | 117 | SNP |
| **C:35131** | 101 | SNP |
| **C:35192** | 119 | SNP |
| **C:35198** | 77 | SNP |
| **C:6276** | 78 | SNP |
| **C:3531** | 119 | SNP |
| **C:3532** | 36 | CHH |
| **C:3533** | 130 | CHG |
| **C:634** | 6 | SNP |
| **C:3537** | 181 | SNP |
| **C:35388** | 81 | SNP |
| **C:354** | 22 | SNP |
| **C:35409** | 114 | SNP |
| **C:3545** | 38 | SNP |
| **C:35534** | 65 | CG |
| **C:35581** | 95 | SNP |
| **C:35609** | 13 | SNP |
| **C:35619** | 11 | SNP |
| **C:35649** | 108 | SNP |

| | | |
|---|---|---|
| **C:35650** | 28 | SNP |
| **C:35650** | 80 | SNP |
| **C:6542** | 74 | SNP |
| **C:6542** | 100 | SNP |
| **C:6542** | 71 | SNP |
| **C:35808** | 106 | SNP |
| **C:6565** | 126 | CG |
| **C:6565** | 100 | CG |
| **C:6565** | 99 | CG |
| **C:7597** | 16 | SNP |
| **C:6565** | 93 | CG |
| **C:3593** | 45 | SNP |
| **C:6571** | 57 | SNP |
| **C:6578** | 193 | SNP |
| **C:36103** | 54 | SNP |
| **C:6582** | 136 | SNP |
| **C:3616** | 88 | SNP |
| **C:3618** | 40 | SNP |
| **C:362** | 7 | SNP |
| **C:362** | 52 | SNP |
| **C:362** | 167 | SNP |
| **C:3622** | 185 | SNP |
| **C:66** | 34 | SNP |
| **C:363** | 184 | SNP |
| **C:3631** | 7 | SNP |
| **C:3672** | 35 | SNP |
| **C:3672** | 75 | SNP |
| **C:6664** | 47 | SNP |
| **C:368** | 128 | SNP |
| **C:3691** | 30 | SNP |
| **C:3692** | 24 | CHH |
| **C:3694** | 17 | SNP |

| | | |
|---|---|---|
| **C:3698** | 66 | SNP |
| **C:7912** | 87 | SNP |
| **C:4516** | 28 | SNP |
| **C:3719** | 9 | SNP |
| **C:372** | 94 | CHH |
| **C:3722** | 54 | SNP |
| **C:3722** | 69 | SNP |
| **C:3722** | 145 | SNP |
| **C:3722** | 173 | SNP |
| **C:3727** | 166 | SNP |
| **C:3727** | 67 | CHG |
| **C:3729** | 115 | SNP |
| **C:6855** | 179 | CG |
| **C:3740** | 34 | SNP |
| **C:6893** | 21 | CHH |
| **C:3769** | 13 | SNP |
| **C:3769** | 67 | SNP |
| **C:3769** | 88 | SNP |
| **C:6913** | 72 | SNP |
| **C:378** | 7 | SNP |
| **C:378** | 185 | SNP |
| **C:3783** | 24 | SNP |
| **C:379** | 176 | SNP |
| **C:380** | 48 | SNP |
| **C:381** | 26 | SNP |
| **C:381** | 88 | SNP |
| **C:3850** | 8 | SNP |
| **C:3850** | 11 | SNP |
| **C:3850** | 109 | SNP |
| **C:3854** | 49 | SNP |
| **C:7** | 133 | CHG |
| **C:3873** | 94 | SNP |

| | | |
|---|---|---|
| **C:3873** | 17 | CHG |
| **C:388** | 61 | SNP |
| **C:39** | 179 | CG |
| **C:3900** | 31 | SNP |
| **C:3900** | 149 | SNP |
| **C:392** | 49 | SNP |
| **C:392** | 110 | SNP |
| **C:393** | 25 | SNP |
| **C:393** | 76 | SNP |
| **C:3932** | 49 | SNP |
| **C:3932** | 64 | SNP |
| **C:3949** | 77 | SNP |
| **C:3950** | 79 | SNP |
| **C:3950** | 121 | SNP |
| **C:3961** | 170 | SNP |
| **C:3962** | 99 | CG |
| **C:3962** | 94 | SNP |
| **C:3979** | 57 | SNP |
| **C:3981** | 51 | SNP |
| **C:400** | 148 | SNP |
| **C:400** | 181 | SNP |
| **C:4008** | 82 | SNP |
| **C:4916** | 8 | SNP |
| **C:4024** | 9 | SNP |
| **C:7319** | 82 | SNP |
| **C:4049** | 16 | CG |
| **C:4049** | 28 | CG |
| **C:4065** | 8 | SNP |
| **C:4085** | 11 | SNP |
| **C:4085** | 138 | SNP |
| **C:4087** | 49 | SNP |
| **C:4113** | 189 | SNP |

| | | |
|---|---|---|
| **C:412** | 50 | SNP |
| **C:412** | 161 | SNP |
| **C:738** | 70 | SNP |
| **C:414** | 196 | SNP |
| **C:415** | 18 | SNP |
| **C:415** | 112 | SNP |
| **C:4155** | 114 | CG |
| **C:416** | 157 | SNP |
| **C:4171** | 168 | SNP |
| **C:418** | 11 | SNP |
| **C:418** | 38 | SNP |
| **C:4198** | 70 | SNP |
| **C:4204** | 15 | SNP |
| **C:4207** | 159 | SNP |
| **C:421** | 70 | SNP |
| **C:4219** | 132 | SNP |
| **C:423** | 80 | SNP |
| **C:4239** | 54 | SNP |
| **C:4252** | 100 | SNP |
| **C:4261** | 39 | SNP |
| **C:4271** | 77 | SNP |
| **C:7500** | 103 | SNP |
| **C:4274** | 151 | SNP |
| **C:4276** | 35 | SNP |
| **C:4287** | 111 | CHH |
| **C:5291** | 20 | SNP |
| **C:431** | 141 | SNP |
| **C:431** | 193 | SNP |
| **C:7722** | 124 | SNP |
| **C:436** | 52 | SNP |
| **C:436** | 138 | SNP |
| **C:4361** | 11 | SNP |

| | | |
|---|---|---|
| **C:4361** | 75 | SNP |
| **C:7773** | 48 | SNP |
| **C:4413** | 135 | SNP |
| **C:7778** | 124 | SNP |
| **C:442** | 132 | SNP |
| **C:778** | 159 | CHH |
| **C:4430** | 183 | SNP |
| **C:4466** | 68 | SNP |
| **C:4472** | 23 | SNP |
| **C:4472** | 137 | SNP |
| **C:79** | 139 | SNP |
| **C:4495** | 45 | SNP |
| **C:79** | 141 | SNP |
| **C:5398** | 11 | SNP |
| **C:541** | 87 | SNP |
| **C:4518** | 78 | SNP |
| **C:4521** | 168 | SNP |
| **C:4524** | 154 | SNP |
| **C:4531** | 12 | SNP |
| **C:8028** | 144 | SNP |
| **C:8029** | 65 | SNP |
| **C:8029** | 116 | CHG |
| **C:8040** | 45 | SNP |
| **C:4606** | 74 | SNP |
| **C:4608** | 65 | SNP |
| **C:461** | 183 | CHH |
| **C:8083** | 45 | SNP |
| **C:8092** | 85 | SNP |
| **C:4629** | 7 | SNP |
| **C:4629** | 104 | SNP |
| **C:4630** | 116 | SNP |
| **C:4630** | 26 | SNP |

| | | |
|---|---|---|
| **C:4630** | 74 | SNP |
| **C:4630** | 109 | SNP |
| **C:464** | 73 | SNP |
| **C:464** | 87 | SNP |
| **C:4649** | 149 | SNP |
| **C:467** | 58 | SNP |
| **C:4685** | 136 | SNP |
| **C:4687** | 49 | SNP |
| **C:8206** | 6 | SNP |
| **C:8230** | 111 | SNP |
| **C:4710** | 170 | CHH |
| **C:4726** | 166 | CHH |
| **C:4763** | 27 | SNP |
| **C:4780** | 56 | CHG |
| **C:591** | 152 | SNP |
| **C:5942** | 135 | SNP |
| **C:4787** | 29 | SNP |
| **C:4791** | 80 | CHG |
| **C:48** | 131 | SNP |
| **C:8365** | 71 | SNP |
| **C:5980** | 38 | SNP |
| **C:4821** | 87 | SNP |
| **C:4824** | 136 | SNP |
| **C:4830** | 38 | SNP |
| **C:4830** | 83 | SNP |
| **C:8436** | 194 | SNP |
| **C:849** | 63 | SNP |
| **C:4860** | 153 | SNP |
| **C:4867** | 101 | SNP |
| **C:8574** | 16 | SNP |
| **C:8579** | 169 | SNP |
| **C:6039** | 187 | SNP |

| | | |
|---|---|---|
| **C:8602** | 182 | SNP |
| **C:4913** | 88 | SNP |
| **C:4913** | 110 | SNP |
| **C:7263** | 125 | SNP |
| **C:4916** | 80 | SNP |
| **C:4946** | 23 | SNP |
| **C:4969** | 130 | SNP |
| **C:4969** | 133 | SNP |
| **C:4975** | 56 | SNP |
| **C:4978** | 46 | CHG |
| **C:50** | 92 | SNP |
| **C:5004** | 165 | SNP |
| **C:5018** | 75 | SNP |
| **C:5047** | 111 | SNP |
| **C:505** | 46 | SNP |
| **C:5055** | 69 | SNP |
| **C:8735** | 23 | SNP |
| **C:629** | 113 | SNP |
| **C:507** | 138 | SNP |
| **C:507** | 185 | SNP |
| **C:5082** | 111 | CG |
| **C:8765** | 86 | SNP |
| **C:5105** | 129 | CG |
| **C:513** | 153 | SNP |
| **C:5138** | 173 | CHH |
| **C:5138** | 109 | CHG |
| **C:5150** | 127 | SNP |
| **C:5151** | 114 | SNP |
| **C:5151** | 150 | SNP |
| **C:5151** | 127 | SNP |
| **C:5187** | 66 | SNP |
| **C:5193** | 189 | SNP |

| | | |
|---|---|---|
| **C:5198** | 149 | SNP |
| **C:5198** | 63 | SNP |
| **C:5198** | 83 | SNP |
| **C:52** | 174 | SNP |
| **C:887** | 108 | SNP |
| **C:887** | 197 | SNP |
| **C:8880** | 56 | CHG |
| **C:525** | 116 | SNP |
| **C:5262** | 60 | SNP |
| **C:755** | 146 | SNP |
| **C:5291** | 176 | CG |
| **C:5292** | 72 | SNP |
| **C:53** | 42 | SNP |
| **C:5322** | 105 | CHH |
| **C:5326** | 126 | CG |
| **C:5326** | 173 | CG |
| **C:5326** | 30 | CG |
| **C:5326** | 25 | CG |
| **C:5326** | 130 | CG |
| **C:5326** | 163 | CHG |
| **C:5326** | 116 | CHG |
| **C:535** | 41 | SNP |
| **C:9192** | 76 | SNP |
| **C:535** | 188 | SNP |
| **C:921** | 15 | SNP |
| **C:5355** | 73 | SNP |
| **C:5355** | 96 | SNP |
| **C:5371** | 194 | SNP |
| **C:5386** | 17 | SNP |
| **C:5387** | 98 | SNP |
| **C:5387** | 142 | SNP |
| **C:539** | 171 | SNP |

| | | |
|---|---|---|
| **C:5390** | 75 | SNP |
| **C:9383** | 49 | SNP |
| **C:6692** | 27 | SNP |
| **C:9392** | 74 | SNP |
| **C:5449** | 32 | CG |
| **C:5461** | 86 | SNP |
| **C:5462** | 64 | CG |
| **C:55** | 16 | CG |
| **C:55** | 18 | CG |
| **C:5510** | 191 | SNP |
| **C:553** | 107 | SNP |
| **C:951** | 71 | SNP |
| **C:9513** | 186 | SNP |
| **C:9518** | 57 | SNP |
| **C:554** | 90 | SNP |
| **C:554** | 93 | SNP |
| **C:554** | 109 | SNP |
| **C:5549** | 66 | SNP |
| **C:9629** | 59 | SNP |
| **C:9671** | 28 | SNP |
| **C:9675** | 40 | SNP |
| **C:8116** | 80 | SNP |
| **C:563** | 185 | SNP |
| **C:565** | 63 | SNP |
| **C:5678** | 72 | SNP |
| **C:57** | 188 | SNP |
| **C:5729** | 111 | SNP |
| **C:5743** | 171 | SNP |
| **C:5745** | 68 | SNP |
| **C:5750** | 31 | SNP |
| **C:5781** | 28 | SNP |
| **C:582** | 142 | SNP |

| | | |
|---|---|---|
| **C:583** | 119 | SNP |
| **C:583** | 124 | SNP |
| **C:5843** | 112 | CG |
| **C:5847** | 10 | SNP |
| **C:5906** | 121 | SNP |
| **C:5906** | 157 | SNP |
| **C:7037** | 71 | SNP |
| **C:9886** | 110 | SNP |
| **C:5959** | 58 | SNP |
| **C:9894** | 86 | SNP |
| **C:9894** | 21 | SNP |
| **C:9900** | 37 | SNP |
| **C:7086** | 39 | SNP |
| **C:60** | 41 | SNP |
| **C:60** | 133 | SNP |
| **C:60** | 148 | SNP |
| **C:60** | 167 | CG |
| **C:6005** | 54 | SNP |
| **C:601** | 96 | SNP |
| **C:6013** | 93 | SNP |
| **C:6016** | 150 | SNP |
| **C:603** | 30 | SNP |
| **C:6039** | 63 | SNP |
| **C:722** | 147 | SNP |
| **C:7228** | 80 | SNP |
| **C:725** | 167 | SNP |
| **C:8636** | 60 | SNP |
| **C:7296** | 55 | SNP |
| **C:610** | 58 | SNP |
| **C:87** | 93 | CG |
| **C:6134** | 35 | SNP |
| **C:6152** | 83 | SNP |

| | | |
|---|---|---|
| **C:618** | 160 | SNP |
| **C:62** | 50 | SNP |
| **C:62** | 166 | SNP |
| **C:622** | 147 | CG |
| **C:7340** | 103 | SNP |
| **C:623** | 125 | SNP |
| **C:624** | 160 | SNP |
| **C:626** | 31 | SNP |
| **C:6622** | 11 | SNP |
| **C:63** | 150 | CHH |
| **C:739** | 53 | SNP |
| **C:739** | 78 | SNP |
| **C:635** | 83 | SNP |
| **C:6358** | 48 | SNP |
| **C:638** | 166 | SNP |
| **C:6387** | 42 | SNP |
| **C:6403** | 100 | SNP |
| **C:6425** | 19 | SNP |
| **C:644** | 95 | SNP |
| **C:6471** | 63 | SNP |
| **C:6519** | 114 | CG |
| **C:653** | 112 | SNP |
| **C:653** | 129 | SNP |
| **C:7472** | 77 | SNP |
| **C:7482** | 53 | SNP |
| **C:749** | 46 | SNP |
| **C:6542** | 135 | SNP |
| **C:6549** | 126 | SNP |
| **C:7503** | 187 | CHG |
| **C:7510** | 41 | SNP |
| **C:8899** | 80 | SNP |
| **C:8908** | 75 | SNP |

| | | |
|---|---|---|
| **C:7597** | 41 | SNP |
| **C:6571** | 48 | SNP |
| **C:760** | 14 | SNP |
| **C:8945** | 146 | SNP |
| **C:6578** | 157 | SNP |
| **C:7723** | 116 | SNP |
| **C:6585** | 10 | SNP |
| **C:659** | 196 | SNP |
| **C:659** | 152 | CHH |
| **C:659** | 132 | CG |
| **C:6590** | 36 | SNP |
| **C:6593** | 92 | SNP |
| **C:6996** | 8 | SNP |
| **C:6607** | 79 | SNP |
| **C:9835** | 75 | SNP |
| **C:6635** | 17 | SNP |
| **C:6649** | 9 | SNP |
| **C:665** | 141 | SNP |
| **C:7791** | 29 | SNP |
| **C:6667** | 48 | SNP |
| **C:788** | 10 | SNP |
| **C:6680** | 130 | SNP |
| **C:6686** | 80 | SNP |
| **C:795** | 12 | SNP |
| **C:7974** | 16 | SNP |
| **C:6725** | 79 | SNP |
| **C:8006** | 32 | SNP |
| **C:678** | 191 | CHH |
| **C:6818** | 105 | SNP |
| **C:6834** | 40 | CG |
| **C:6837** | 52 | SNP |
| **C:6844** | 43 | CHG |

| | | |
|---|---|---|
| **C:8083** | 22 | SNP |
| **C:6871** | 115 | SNP |
| **C:9577** | 146 | SNP |
| **C:6893** | 57 | CG |
| **C:69** | 67 | SNP |
| **C:690** | 136 | SNP |
| **C:7319** | 81 | SNP |
| **C:6925** | 122 | CHG |
| **C:6953** | 12 | SNP |
| **C:6953** | 135 | SNP |
| **C:696** | 41 | SNP |
| **C:6970** | 61 | SNP |
| **C:6989** | 145 | SNP |
| **C:699** | 131 | SNP |
| **C:7361** | 26 | SNP |
| **C:7** | 88 | SNP |
| **C:8731** | 76 | CHH |
| **C:701** | 6 | SNP |
| **C:7027** | 44 | SNP |
| **C:8257** | 68 | SNP |
| **C:832** | 54 | SNP |
| **C:739** | 123 | SNP |
| **C:989** | 174 | SNP |
| **C:7398** | 49 | SNP |
| **C:7410** | 77 | SNP |
| **C:742** | 157 | CG |
| **C:7086** | 81 | CG |
| **C:7087** | 8 | CG |
| **C:7088** | 68 | CG |
| **C:709** | 70 | SNP |
| **C:709** | 70 | CG |
| **C:709** | 71 | CG |

| | | |
|---|---|---|
| **C:71** | 89 | SNP |
| **C:7124** | 56 | SNP |
| **C:854** | 22 | SNP |
| **C:7185** | 166 | SNP |
| **C:8040** | 90 | SNP |
| **C:722** | 161 | SNP |
| **C:8895** | 58 | SNP |
| **C:8602** | 152 | CG |
| **C:8098** | 142 | SNP |
| **C:8664** | 143 | SNP |
| **C:7597** | 49 | SNP |
| **C:8910** | 46 | SNP |
| **C:7319** | 98 | SNP |
| **C:7319** | 111 | SNP |
| **C:7324** | 54 | SNP |
| **C:7326** | 109 | CG |
| **C:7333** | 77 | SNP |
| **C:7335** | 135 | CHG |
| **C:8707** | 112 | SNP |
| **C:7773** | 45 | SNP |
| **C:7361** | 75 | SNP |
| **C:7773** | 115 | SNP |
| **C:9209** | 95 | SNP |
| **C:7385** | 133 | SNP |
| **C:8742** | 14 | SNP |
| **C:8742** | 197 | SNP |
| **C:739** | 119 | SNP |
| **C:7799** | 169 | SNP |
| **C:7393** | 79 | SNP |
| **C:79** | 140 | SNP |
| **C:8345** | 103 | SNP |
| **C:8365** | 62 | SNP |

| | | |
|---|---|---|
| **C:7445** | 32 | SNP |
| **C:7445** | 64 | SNP |
| **C:7445** | 16 | CG |
| **C:7472** | 117 | SNP |
| **C:7472** | 17 | SNP |
| **C:884** | 47 | SNP |
| **C:884** | 36 | SNP |
| **C:8848** | 37 | SNP |
| **C:750** | 179 | CHH |
| **C:7732** | 68 | SNP |
| **C:9553** | 103 | SNP |
| **C:9557** | 140 | CG |
| **C:817** | 110 | SNP |
| **C:8182** | 109 | SNP |
| **C:891** | 115 | SNP |
| **C:8109** | 49 | SNP |
| **C:87** | 85 | CG |
| **C:8116** | 112 | SNP |
| **C:9018** | 31 | SNP |
| **C:773** | 70 | SNP |
| **C:773** | 84 | SNP |
| **C:9341** | 47 | SNP |
| **C:7744** | 178 | CG |
| **C:7773** | 72 | SNP |
| **C:8794** | 79 | SNP |
| **C:9430** | 13 | CHG |
| **C:8183** | 82 | SNP |
| **C:9833** | 109 | SNP |
| **C:778** | 129 | SNP |
| **C:929** | 34 | SNP |
| **C:7781** | 188 | SNP |
| **C:9326** | 81 | SNP |

| | | |
|---|---|---|
| **C:988** | 134 | SNP |
| **C:934** | 162 | SNP |
| **C:8040** | 70 | SNP |
| **C:8340** | 132 | CHH |
| **C:8775** | 136 | SNP |
| **C:8159** | 109 | SNP |
| **C:8707** | 172 | SNP |
| **C:9448** | 164 | SNP |
| **C:7984** | 177 | CHH |
| **C:80** | 90 | SNP |
| **C:949** | 28 | SNP |
| **C:949** | 173 | SNP |
| **C:9504** | 190 | SNP |
| **C:9507** | 104 | CG |
| **C:8533** | 55 | SNP |
| **C:8148** | 123 | SNP |
| **C:8152** | 65 | CG |
| **C:8159** | 100 | SNP |
| **C:9361** | 107 | SNP |
| **C:914** | 45 | CHG |
| **C:8726** | 56 | CG |
| **C:8100** | 94 | SNP |
| **C:8387** | 107 | CHH |
| **C:891** | 131 | SNP |
| **C:8239** | 96 | SNP |
| **C:8117** | 80 | CHH |
| **C:8148** | 9 | SNP |
| **C:933** | 18 | SNP |
| **C:8775** | 159 | SNP |
| **C:8775** | 71 | SNP |
| **C:8707** | 130 | SNP |
| **C:9893** | 83 | SNP |

| | | |
|---|---|---|
| **C:881** | 53 | SNP |
| **C:873** | 40 | SNP |
| **C:960** | 18 | SNP |
| **C:984** | 54 | SNP |
| **C:8407** | 64 | SNP |
| **C:8268** | 116 | SNP |
| **C:8765** | 17 | SNP |
| **C:87** | 87 | CG |
| **C:8700** | 140 | CHG |
| **C:8700** | 176 | CG |
| **C:906** | 45 | SNP |
| **C:9069** | 79 | SNP |
| **C:918** | 18 | CG |
| **C:9900** | 60 | SNP |
| **C:982** | 21 | SNP |
| **C:84** | 19 | SNP |
| **C:883** | 122 | SNP |
| **C:8426** | 6 | SNP |
| **C:884** | 83 | SNP |
| **C:87** | 88 | CG |
| **C:9742** | 176 | CHH |
| **C:902** | 76 | SNP |
| **C:9749** | 10 | SNP |
| **C:9751** | 19 | CG |
| **C:9772** | 160 | SNP |
| **C:9187** | 31 | SNP |
| **C:8810** | 144 | SNP |
| **C:8815** | 110 | SNP |
| **C:9738** | 125 | SNP |
| **C:9263** | 39 | SNP |
| **C:87** | 94 | CG |
| **C:9984** | 69 | SNP |

| | | |
|---|---|---|
| **C:9978** | 104 | SNP |
| **C:976** | 29 | CG |
| **C:98** | 186 | SNP |
| **C:9815** | 119 | SNP |
| **C:8836** | 85 | SNP |
| **C:9978** | 107 | SNP |
| **C:8831** | 105 | SNP |
| **C:984** | 96 | SNP |
| **C:9488** | 23 | SNP |
| **C:9967** | 161 | SNP |
| **C:9448** | 112 | SNP |
