## Supplementary figures and images for "Reduced representation characterization of genetic and epigenetic differentiation to oil pollution in the foundation plant *Spartina alterniflora*"

### Figure S1

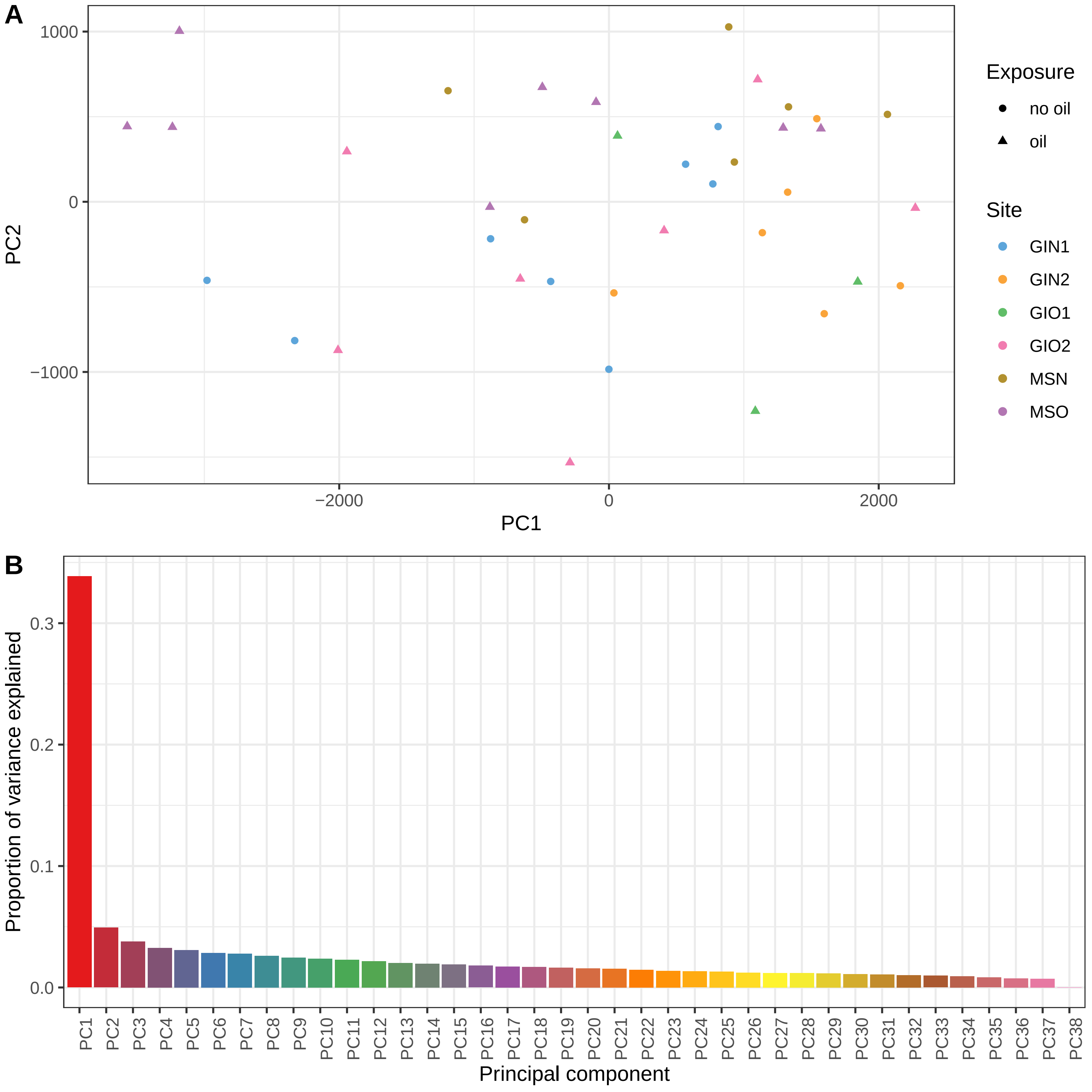

### Figure S2

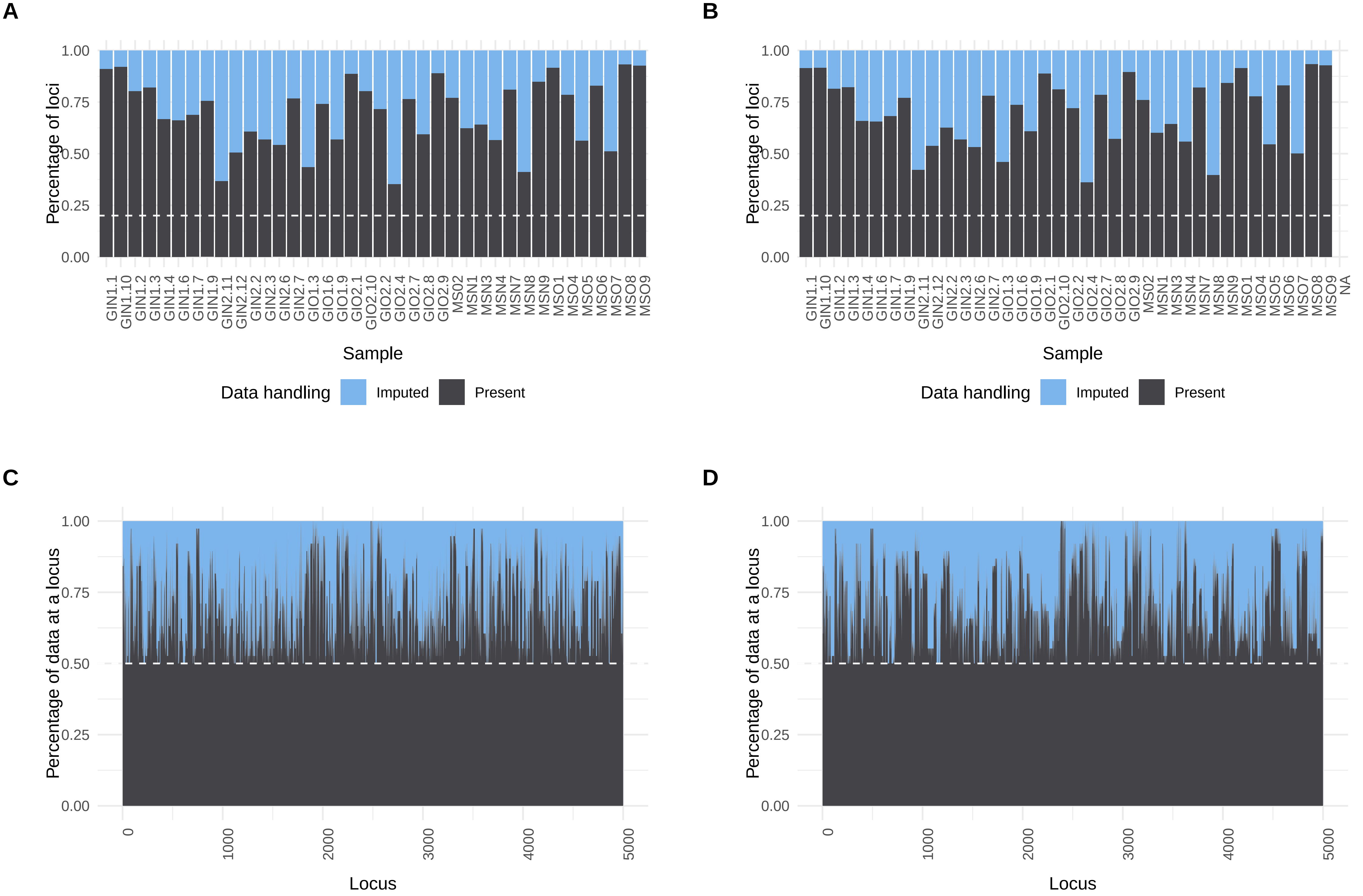
